# A PUF hub drives self-renewal in *C. elegans* germline stem cells

**DOI:** 10.1101/785923

**Authors:** Kimberly A. Haupt, Kimberley T. Law, Amy L. Enright, Charlotte R. Kanzler, Heaji Shin, Marvin Wickens, Judith Kimble

## Abstract

Stem cell regulation relies on extrinsic signaling from a niche plus intrinsic factors that respond and drive self-renewal within stem cells. A *priori,* loss of niche signaling and loss of the intrinsic self-renewal factors might be expected to have equivalent stem cell defects. Yet this simple prediction has not been borne out for most stem cells, including *C. elegans* germline stem cells (GSCs). The central regulators of *C. elegans* GSCs include extrinsically-acting GLP-1/Notch signaling from the niche, intrinsically-acting RNA binding proteins in the PUF family, termed FBF-1 and FBF-2 (collectively FBF), and intrinsically-acting PUF partner proteins that are direct Notch targets. Abrogation of either GLP-1/Notch signaling or its targets yields an earlier and more severe GSC defect than loss of FBF-1 and FBF-2, suggesting that additional intrinsic regulators must exist. Here, we report that those missing regulators are two additional PUF proteins, PUF-3 and PUF-11. Remarkably, an *fbf-1 fbf-2; puf-3 puf-11* quadruple null mutant has a GSC defect virtually identical to that of a *glp-1/*Notch null mutant. PUF-3 and PUF-11 both affect GSC maintenance; both are expressed in GSCs; and epistasis experiments place them at the same position as FBF within the network. Therefore, action of PUF-3 and PUF-11 explains the milder GSC defect in *fbf-1 fbf-2* mutants. We conclude that a “PUF hub”, comprising four PUF proteins and two PUF partners, constitutes the intrinsic self-renewal node of the *C. elegans* GSC RNA regulatory network. Discovery of this hub underscores the significance of PUF RNA-binding proteins as key regulators of stem cell maintenance.

## INTRODUCTION

Stem cell regulatory networks govern the balance between self-renewal and differentiation. Transcription factors are well established stem cell regulators (e.g. BOYER *et al*. 2005), as are RNA-binding proteins (e.g. WICKENS *et al*. 2002; YE AND BLELLOCH 2014; GROSS-THEBING *et al*. 2017). Indeed, the PUF (for <u>Pu</u>milio and <u>F</u>BF) family of RNA-binding proteins promote stem cell self-renewal in multiple tissues across animal phylogeny from planaria to mammals (LIN AND SPRADLING 1997; FORBES AND LEHMANN 1998; CRITTENDEN *et al*. 2002; SALVETTI *et al*. 2005; NAUDIN *et al*. 2017; ZHANG *et al*. 2017). Genome-wide studies reveal that PUF proteins bind hundreds of RNAs (HAFNER *et al*. 2010; KERSHNER AND KIMBLE 2010; PRASAD *et al*. 2016; PORTER *et al*. 2019), consistent with a central role in the stem cell regulatory network (KERSHNER *et al*. 2013). Yet the full significance of PUF proteins in self-renewal has been unclear. Are PUF proteins the primary stem-cell intrinsic self-renewal regulators? Or do they play only a supporting role? This question has been difficult to tackle in organisms where only one or two PUF proteins are responsible for many diverse processes, including embryogenesis, germ cell development and neural activities. Here we address the question in nematodes, where the number of genes encoding PUF proteins has expanded during evolution, yielding functional specialization among family members.

We focus here on the role of PUF proteins in regulating self-renewal in the *C. elegans* germline. Germline stem cells (GSCs) expand the germline tissue from two cells at hatching to ∼2000 cells in adults; they replenish the tissue as germ cells are lost to gametogenesis during reproduction (KIMBLE AND WHITE 1981; CRITTENDEN *et al*. 2006); and they regenerate the tissue upon feeding after starvation (ANGELO AND VAN GILST 2009; SEIDEL AND KIMBLE 2011). Key regulators of GSC self-renewal include GLP-1/Notch signaling from the niche; its transcriptional targets, *lst-1* and *sygl-1;* and two PUF proteins, FBF-1 and FBF-2 (collectively FBF) (Figure 1A) (AUSTIN AND KIMBLE 1987; CRITTENDEN *et al*. 2002; KERSHNER *et al*. 2014).

**Figure 1:**
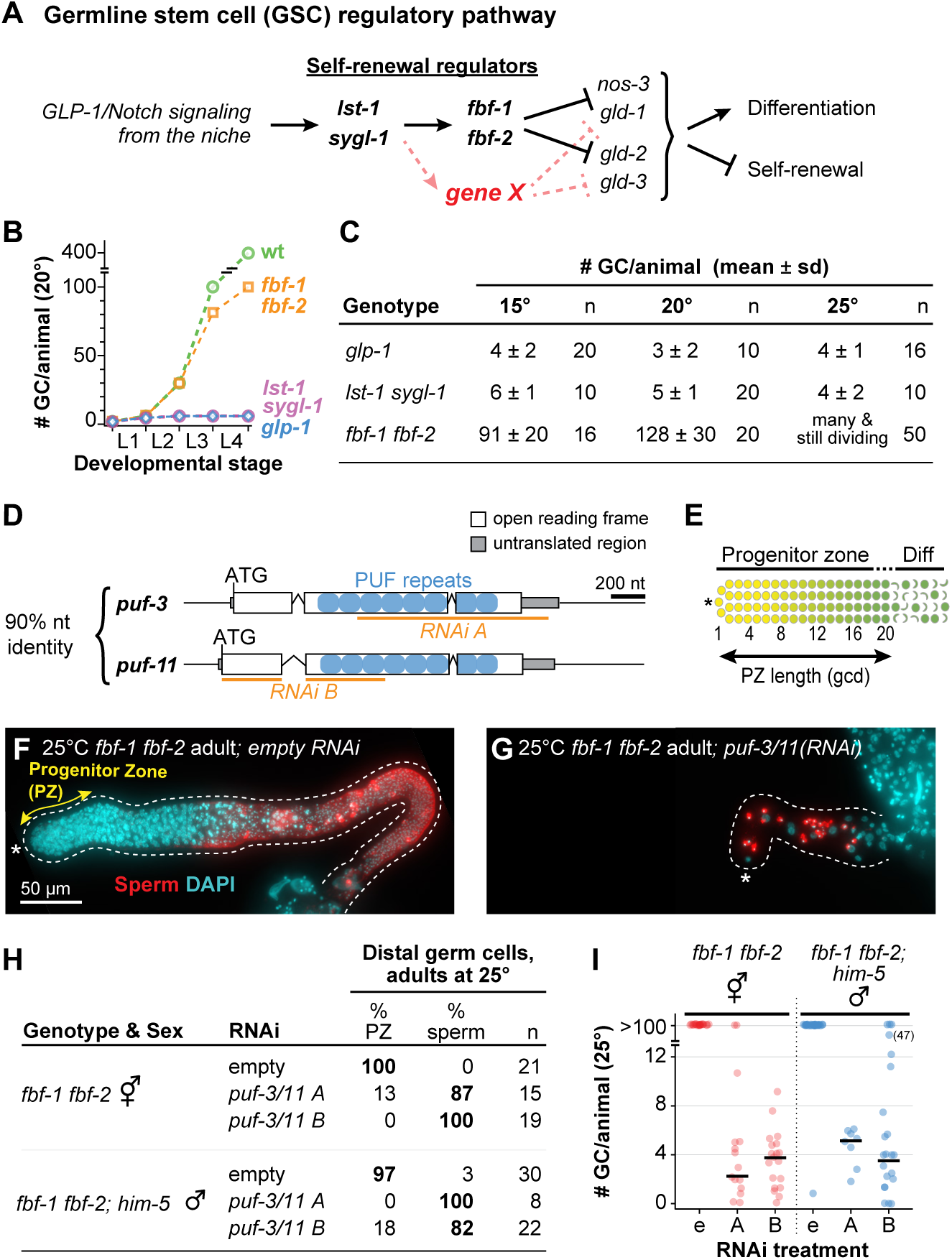
*puf-3* and *puf-11* are putative missing GSC regulators. **A.** Diagram illustrating functional relationships between key genes that regulate stem cell self-renewal and differentiation in *C. elegans* germline stem cells (GSCs). Gene X (red) represents a missing factor that likely functions in parallel to *fbf-1* and *fbf-2*. **B.** Total number of germ cells (GC) per animal at 20° in wild-type (wt) and mutants of GSC regulators: *fbf-1 fbf-2*, *lst-1 sygl-1* and *glp-1* mutants (AUSTIN AND KIMBLE 1987; CRITTENDEN *et al*. 2002; KIMBLE AND CRITTENDEN 2005; CRITTENDEN *et al*. 2006; KERSHNER *et al*. 2014). X-axis: L1-L4, larval stages of development; ticks mark molts between stages. Alleles used here and throughout this work are *fbf-1(ok91), fbf-2(q704), glp-1(q46), lst-1(ok814)* and *sygl-1(tm5040)*. **C.** Total GC number produced per animal. Animals of each strain were staged to 36 hours (15°), 24 hours (20°) or 18 hours (25°) past mid-L4 for this experiment. In germlines where distal germ cells had differentiated to mature sperm, GC number was determined by counting sperm number and dividing by four. GC number were not counted in larger germlines that retained a progenitor zone; these were scored as “many & still dividing”. Alleles are identical to Figure 1B except for *lst-1(q869)* and *sygl-1(q828).* n, number animals scored. **D.** Schematic of *puf-3* and *puf-11* loci, which share 90% nucleotide (nt) identity (HUBSTENBERGER *et al*. 2012): exons (white boxes), introns (peaked lines), untranslated region (gray boxes), sequence encoding individual PUF repeats (blue circles), sequence targeted by RNAi clone (orange line). **E.** Diagram of distal gonad with the progenitor zone (PZ) and germ cells that have entered early meiotic prophase (Diff). Distal end (asterisk); GSCs (yellow); GSC daughters primed for differentiation and transitioning toward entry into meiotic prophase, including those in meiotic S-phase (graded yellow to green); early meiotic prophase (green, crescent-shaped). **F,G.** Representative images of extruded gonads from 25° *fbf-1 fbf-2* adult hermaphrodites staged to 18 hours after L4 on either empty vector control (F) or *puf-3/11* RNAi clone B (G). Gonads are immunostained with a sperm marker SP56 (red) and DAPI (cyan). Images are Z-projections of several fluorescent images obtained on a compound microscope. A dotted line delineates the gonad boundary and an asterisk marks the distal end. Double headed arrow (yellow) indicates the PZ. Scale bar in F applies to both images. **H.** State of distal germ cells of *fbf-1 fbf-2* hermaphrodites and *fbf-1 fbf-2; him-5* males, raised at 25° on either empty or *puf-3/11* RNAi and staged to adulthood, 18 hours after L4. States were determined by DAPI-stained chromosomal morphology, scoring for a PZ or mature sperm in the distal germline. *him-5(e1490)* increased the frequency of male progeny (HODGKIN *et al*. 1979). **I.** Quantitation of total GC per animal from experiment in (H) by counting sperm number and dividing by four. In germlines with a PZ, GC numbers were not counted but appeared comparable to previous reports of >100 (MERRITT AND SEYDOUX 2010; SHIN *et al*. 2017). RNAi treatment on x-axis: empty vector (e), *puf-3/11* RNAi clone A (A) or *puf-3/11* RNAi clone B (B). Individual data points are plotted as circles; middle line is median value.

While FBF-1 and FBF-2 are crucial in GSC control, they cannot account for all the effects of niche signaling. Removal of either the GLP-1/Notch receptor or both *lst-1* and *sygl-1* has an early and severe germline proliferation (Glp) defect in both sexes: the two GSCs at hatching divide once or twice and then differentiate prematurely as sperm (AUSTIN AND KIMBLE 1987; KERSHNER *et al*. 2014). Moreover, GLP-1/Notch and its targets drive self-renewal throughout larval development and in adults (AUSTIN AND KIMBLE 1987; KERSHNER *et al*. 2014). By contrast, GSCs in the *fbf-1 fbf-*2 double mutant are lost to differentiation much later, at the fourth larval stage (L4s) (CRITTENDEN *et al*. 2002). Indeed, this delayed L4 GSC defect occurs in mutants raised at 15° and 20°, but at 25°, the effect is even further delayed with GSCs continuing to divide in adults and persisting in an undifferentiated state (MERRITT *et al*. 2008; SHIN *et al*. 2017; this work). We designate the less severe germline phenotype of *fbf-1 fbf-*2 mutants pGlp (“partial Glp”) to distinguish it from the more severe Glp GSC defect seen upon loss of niche signaling. The pGlp defect implies that some other self-renewal factor, “gene X” (Figure 1A), must maintain GSCs in larvae at all temperatures and in adults at 25°.

Here, we report that two additional PUF proteins, PUF-3 and PUF-11, are gene X. When PUF-3 and PUF-11 are removed in an *fbf-1 fbf-2* mutant, the GSC defect is essentially the same as that seen upon loss of niche signaling. Additional results solidify the conclusion that PUF-3 and PUF-11 are intrinsic regulators of GSC self-renewal. We conclude that a “PUF hub”, comprising four PUF proteins and two PUF partners, constitutes the intrinsic self-renewal node of the *C. elegans* GSC RNA regulatory network. Discovery of this hub underscores the significance of PUF RNA-binding proteins as key regulators of stem cell maintenance.

## MATERIALS AND METHODS

### Nematode strains and maintenance

*C. elegans* strains were maintained at 20°, unless specified otherwise, on nematode growth medium (NGM) plates spotted with *E. coli* OP50 (BRENNER 1974). Wild-type was N2 Bristol strain. For a complete list of strains used in this study, see Table S1. *puf-11(gk203683)* is from the Million Mutations Project (THOMPSON *et al*. 2013). We also used the following balancers: *hT2*[*qIs48*] *I;III* (SIEGFRIED AND KIMBLE 2002), *mIn1*[*mIs14dpy-10(e128)*] *II* (EDGLEY AND RIDDLE 2001) and *nT1*[*qIs51*] *IV;V (EDGLEY et al. 2006)*.

Due to incompatibility between *mIn1* and *nT1* balancers, we generated a “pseudo (Ψ)-balancer” to maintain quadruple mutant strains. This *LG II* Ψ-balancer harbors a transgene driving expression of a red fluorescent protein in somatic nuclei, *oxTi564* [*Peft-3::tdTomato::H2B::unc-54 3’UTR + Cbr-unc-119(+)*] (FRØKJÆR-JENSEN *et al*. 2014) plus a closely linked *dpy-10(q1074)* deletion. Quadruple mutants were thus maintained as *fbf-1 fbf-2/* Ψ-balancer [*dpy-10(q1074) oxTi564*] *II*; *puf-3 puf-11 IV/nT1*[*qIs51*] *IV;V*.

### CRISPR/Cas9 genome editing

We used RNA-protein complex CRISPR/Cas9 genome editing with a co-conversion strategy (ARRIBERE *et al*. 2014; PAIX *et al*. 2015) to generate a number of alleles for this work (see Table S2 for details). For each edit, we prepared an injection mix containing: gene-specific crRNAs (10 µM, IDT-Alt-R^TM^); *dpy-10* or *unc-58* co-CRISPR crRNAs (4 µM, IDT-Alt-R^TM^); tracrRNAs (13.6 µM, IDT-Alt-R^TM^); gene specific repair oligo (4 µM); *dpy-10* or *unc-58* repair oligo (1.34 µM); and Cas-9 protein (24.5 µM). Wild-type germlines were injected to make *puf-3(q966), puf-11(q971), puf-3(q1058)* and *puf-11(q1128)*; EG7866 germlines were injected for *dpy-10(q1074)*. F1 progeny of injected hermaphrodites were screened for desired mutations by PCR and Sanger sequencing. Each allele was outcrossed with wild-type at least twice prior to analysis.

### Isolation of *puf-3(q801)*

The *puf-3(q801)* 776 nucleotide deletion allele was isolated from a mutagenized library (gift of Maureen Barr), PCR screened to identify homozygotes and outcrossed against wild-type six times before analysis.

### RNA interference

For RNA interference (RNAi), we identified clones targeting *puf-3, puf-5, puf-6, puf-7, puf-8* and *puf-9* from the Ahringer library (FRASER *et al*. 2000). The library does not include a clone targeting *puf-11*, so we generated one using the Gibson assembly method (GIBSON 2009). Nucleotides 1-800 of the *puf-11* ORF were amplified from *C. elegans* cDNA and cloned into the plasmid L4440 at the *Nco* I site. For all RNAi experiments, the L4440 plasmid lacking a gene of interest (“empty” RNAi) served as a control.

To perform RNAi by feeding (TIMMONS AND FIRE 1998), *E. coli HT115(DE3)* were transformed with RNAi vectors and cultured at 37° overnight in 2xYT media containing 25 μg/μl carbenicillin and 50 μg/μl tetracycline. Bacterial cultures were concentrated and seeded onto NGM plates containing 1mM IPTG, then induced overnight at RT. We then placed mid L4 hermaphrodites on these plates. For experiments shown in Figure 3D,E, we assayed treated animals 48 hours after plating. For all other experiments, treated animals were allowed to lay eggs and their F1 progeny were assayed for defects.

### Analysis and quantification of germ cells in adults

We first staged animals to roughly the same stage of adulthood. Specifically, we picked mid-L4s raised at 15°, 20° or 25° for at least one generation and grew them for an additional 18 hours (25°), 24 hours (20°) or 36 hours (15°) to synchronize adults for analysis. Whole worms were then DAPI stained and imaged by compound microscopy. The presence or absence of a progenitor zone (PZ) was assayed by nuclear morphology of germ cells at the distal end of the gonad: those lacking meiotic nuclear morphology and often possessing M-phase or anaphase nuclei were scored as PZ-positive (see also PZ analysis, below); those having a meiotic prophase nuclear morphology were scored as “meiotic” and PZ-negative; arms where all germ cells had differentiated as mature sperm were scored as “sperm” and PZ-negative. To estimate total germ cell number, we counted mature sperm (which have a distinctive, compact DAPI morphology) using the Multipoint Tool in Fiji/ImageJ (SCHINDELIN *et al*. 2012) and then divided sperm number by four (each germ cell makes four sperm). In some cases, we counted sperm number in one gonadal arm and multiplied that number by two to estimate total sperm number. In cases where all germ cells in an animal had not yet differentiated to sperm, we did not count total germ cells.

### Phenotype analyses: Fertility, brood size, and embryo viability

L4 hermaphrodites were placed onto individual plates at 20°. At 6- to 24-hour intervals, each hermaphrodite was moved to a new plate and the embryos were counted to score for fertility and determine brood size. Several days later, hatched progeny on each plate were counted to determine embryo lethality.

### Progenitor zone (PZ) analysis

From roughly staged adults as described above, gonads were extruded, DAPI stained, and imaged by compound or confocal microscopy. We examined morphology of germline nuclei to determine PZ size according to convention (CRITTENDEN *et al*. 2006; SEIDEL AND KIMBLE 2015) (see also Figure 1E). Briefly, DAPI-staining of nuclei in early meiotic prophase adopts a crescent shape. To count number of germ cells in a PZ, we counted the total number of cells in the distal germline that had not entered early meiotic prophase, using Fiji/ImageJ Cell Counter plugin (SCHINDELIN *et al*. 2012). We also measured the distance to the end of the PZ in germ cell diameters (gcd) from the distal end. To this end, we selected a middle focal plane and counted the number of germline nuclei along each edge of the gonad until the first one with crescent morphology. We averaged the two values from each edge to determine PZ size.

### Quantification of germ cells in larvae

To generate roughly synchronous embryos, we allowed gravid hermaphrodites to lay eggs for 2 hours at 20°. At subsequent timepoints corresponding to early L1, early L2, late L2 and early L4, larvae were harvested and germ cell number quantitated. For early L1s, we scored germ cell number in live animals by DIC microscopy. For all remaining samples, we used whole mount staining with the reduction/oxidation method (FINNEY AND RUVKUN 1990; MILLER AND SHAKES 1995). Briefly, samples were fixed in Ruvkun Fixation Buffer with 1% (v/v) paraformaldehyde for 30 minutes. Disulfide linkage reduction was performed, then samples were incubated in blocking solution (PBS with 1% (w/v) bovine serum albumin, 0.5% (v/v) Triton X-100, 1 mM EDTA) for 40 minutes at RT, followed by overnight incubation at 4° with rabbit α-PGL-1 (1:100, gift from Susan Strome, University of California, Santa Cruz) diluted in blocking solution. Secondary Alexa 555 donkey α-rabbit (1:1000, Thermo Fisher Scientific #A31570) antibody was diluted in blocking solution and incubated with samples for at least one hour along with 1 ng/μl DAPI for DNA visualization. Samples were mounted in Vectashield (Vector Laboratories #H1000) on 2% agarose pads then assayed by fluorescent compound microscopy. Number of germ cells in L2s was determined by counting the total number of PGL-1-postitive cells. For early L4s, we counted the number of PGL-1-positive cells in one gonadal arm then multiplied by two to estimate the total number of germ cells per animal.

### Immunostaining and DAPI staining

We immunostained gonads as described (CRITTENDEN *et al*. 2017) with minor modifications. To extrude gonads, we dissected animals in PBS buffer with 0.1% (v/v) Tween-20 (PBStw) and 0.25 mM levamisole. Gonads were fixed in 3% (w/v) paraformaldehyde diluted in PBStw for 10 minutes, then permeabilized in either 0.2% (v/v) Triton-X diluted in PBStw or ice-cold methanol for 10-15 minutes. Next, gonads were incubated for at least 1 hour in blocking solution (0.5% (w/v) bovine serum albumin diluted in PBStw) and incubated overnight at 4° with primary antibodies diluted in blocking solution as follows: mouse α-V5 (1:1000, SV5-Pk1, Bio-Rad #MCA1360); mouse α-SP56 (1:200, gift from Susan Strome, University of California, Santa Cruz). Secondary antibody Alexa 488 donkey α-mouse (1:1000, Thermo Fisher Scientific #A21202) was diluted in blocking solution and incubated with samples for at least one hour. To visualize DNA, DAPI (4′,6-diamidino-2-phenylindole) was included with the secondary antibody at a final concentration of 1 ng/μl. Samples were mounted in Vectashield (Vector Laboratories #H1000) or ProLong Gold (Thermo Fisher Scientific #P36930) before imaging. All steps were performed at room temperature unless otherwise indicated. Where only DNA visualization was required, we skipped all blocking solution steps and simply incubated samples in PBStw with 1 ng/μl DAPI for 15 minutes prior to mounting.

### Microscopy

Images in Figure 2C,D, 3D,E, 5B-G and Figure S1A,B and S4B-J were taken using a laser scanning Leica TCS SP8 confocal microscope fitted with both photomultiplier and hybrid detectors and run using LAS software version X. A 63x/1.40 CS2 HC Plan Apochromat oil immersion objective was used. All images were taken with 400 Hz scanning speed and 125-200% zoom. To prepare images for figures, Adobe Photoshop was used to equivalently and linearly adjust intensity among images to be compared. Images in Figure 1F,G, 4A-D and S5 were captured using a Hamamatsu ORCA-Flash4.0 cMOS camera on a Zeiss Axioskop compound microscope equipped with 63x 1.4NA Plan Apochromat oil immersion objective. The fluorescent light source was a Lumencore SOLA Light Engine, and Carl Zeiss filter sets 49 and 38 were used for DAPI and Alexa 488 visualization. The acquisition software was Micromanager (EDELSTEIN *et al*. 2010; EDELSTEIN *et al*. 2014). When required, images were combined using the Pairwise Stitching function in Fiji/ImageJ (PREIBISCH *et al*. 2009). To prepare images for figures, Adobe Photoshop was used to adjust intensity equivalently and linearly among images to be compared.

**Figure 2:**
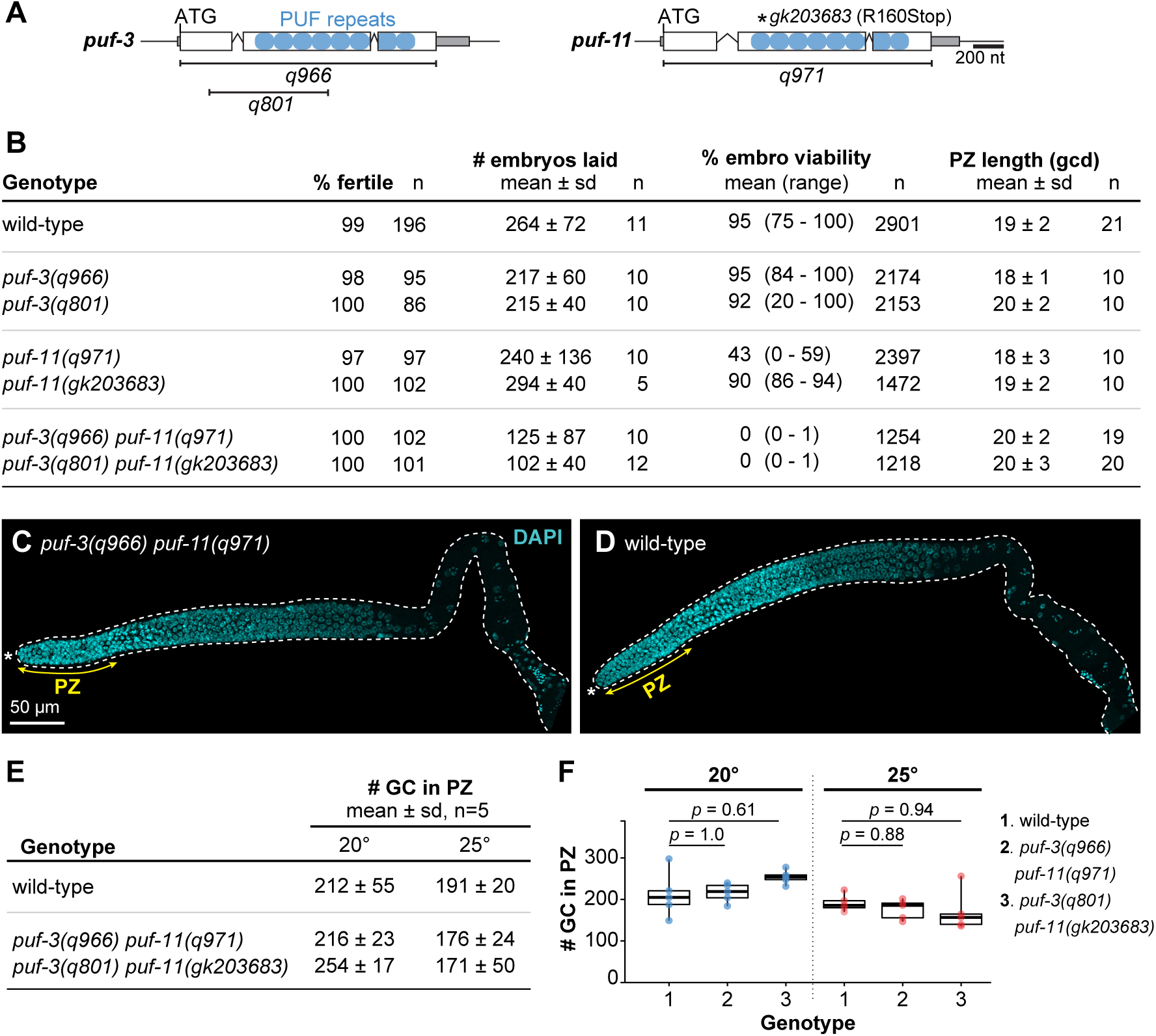
GSC maintenance defects are undetectable in single and double *puf-3* and *puf-11* mutants. **A.** *puf-3* and *puf-11* loci, using conventions as in Figure 1D. Extents of deletion mutants are bracketed below loci and position of the point mutation is marked with an asterisk above. See Figure S2 for sequence details. **B.** Germline-related characteristics of single and double *puf-3* and *puf-11* mutants. See Methods for details about assays and scoring. **C,D.** Representative confocal Z-projections of DAPI-stained gonads extruded from animals staged to 24 hours after L4 at 20°. Extent of progenitor zone (PZ), double headed yellow arrow. Annotation by convention in Figure 1F,G and scale bar in C applies to D. **E,F.** PZ sizes measured in number of germ cells (GC). **F.** PZ sizes showing individual data points as circles; middle line, median; boxes, 25-75% quantile; whiskers, minimum and maximum values. n=5 gonad arms for each sample. *p*-values, mutants were compared to wild-type using Welch’s ANOVA and Games-Howell *post hoc* test.

**Figure 3:**
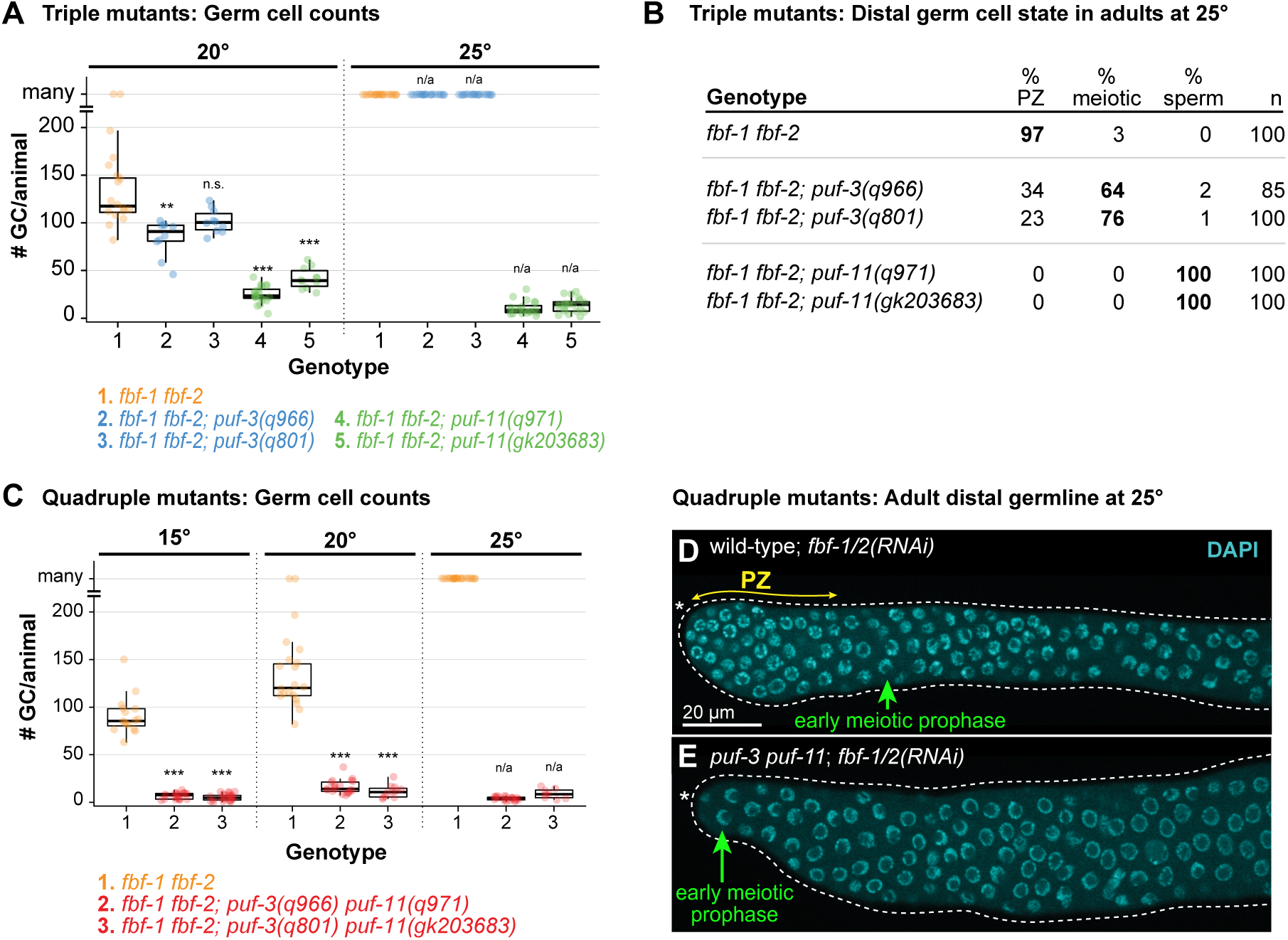
Triple and quadruple mutants reveal *puf-3* and *puf-11* role in GSC maintenance. **A.** Total germ cell (GC) number per animal in *fbf-1 fbf-2; puf* triple mutant adults at 20° and 25°. GC number was counted only in germlines differentiated to the distal end; those with many germ cells and retaining a progenitor zone (PZ) or having meiotic prophase nuclear morphology were scored as “many”. Where distal germ cells had differentiated to mature sperm, GC number was determined by counting sperm and then dividing by four. Individual data points are plotted as circles; middle line, median; boxes, 25-75% quantile; whiskers, minimum and maximum values falling outside the box but within 1.5 times the interquartile range. To determine *p*-values, triple mutants were compared to *fbf-1 fbf-2* control using Welch’s ANOVA and Games-Howell *post-hoc* test; *n.s.*, not significant; ** *p* < 0.01; *** *p* < 0.001; n/a, not applicable. For additional information including mean GC/animal, see Figure S3. **B.** State of distal germ cells in *fbf-1 fbf-2; puf* triple mutants at 25°, assayed in adults at 18 hours past L4. State was scored as described in Figure 1H, with the additional classification of “meiotic” for cells with meiotic prophase chromosomal morphology, but not yet differentiated as sperm. **C.** Graph of germ cell number per animal in *fbf-1 fbf-2; puf-3 puf-11* adults at 15°, 20° and 25°. Germline scoring and graph conventions as in Figure 3A. Individual data points are plotted as circles. *p*-value compared to respective *fbf-1 fbf-2* control was determined using Welch’s ANOVA and Games-Howell *post-hoc* test; *** *p* < 0.001; n/a, not applicable. For additional information including mean GC/animal, see Figure S3. **D,E**. Representative images of DAPI-stained gonads extruded from wild-type (D) or *puf-3(q966) puf-11(q971)* (E) adult hermaphrodites treated with *fbf-1/2* RNAi at 25°. Animals were plated to RNAi as mid-L4s and analyzed 48 hours later for changes in nuclear morphology. Nuclear morphologies indicative of PZ (double headed arrow, yellow) and early meiotic prophase (green) are annotated. Other annotation by convention in Figure 1F,G. Images show single confocal Z-sections and scale bar in D applies to both images.

**Figure 4:**
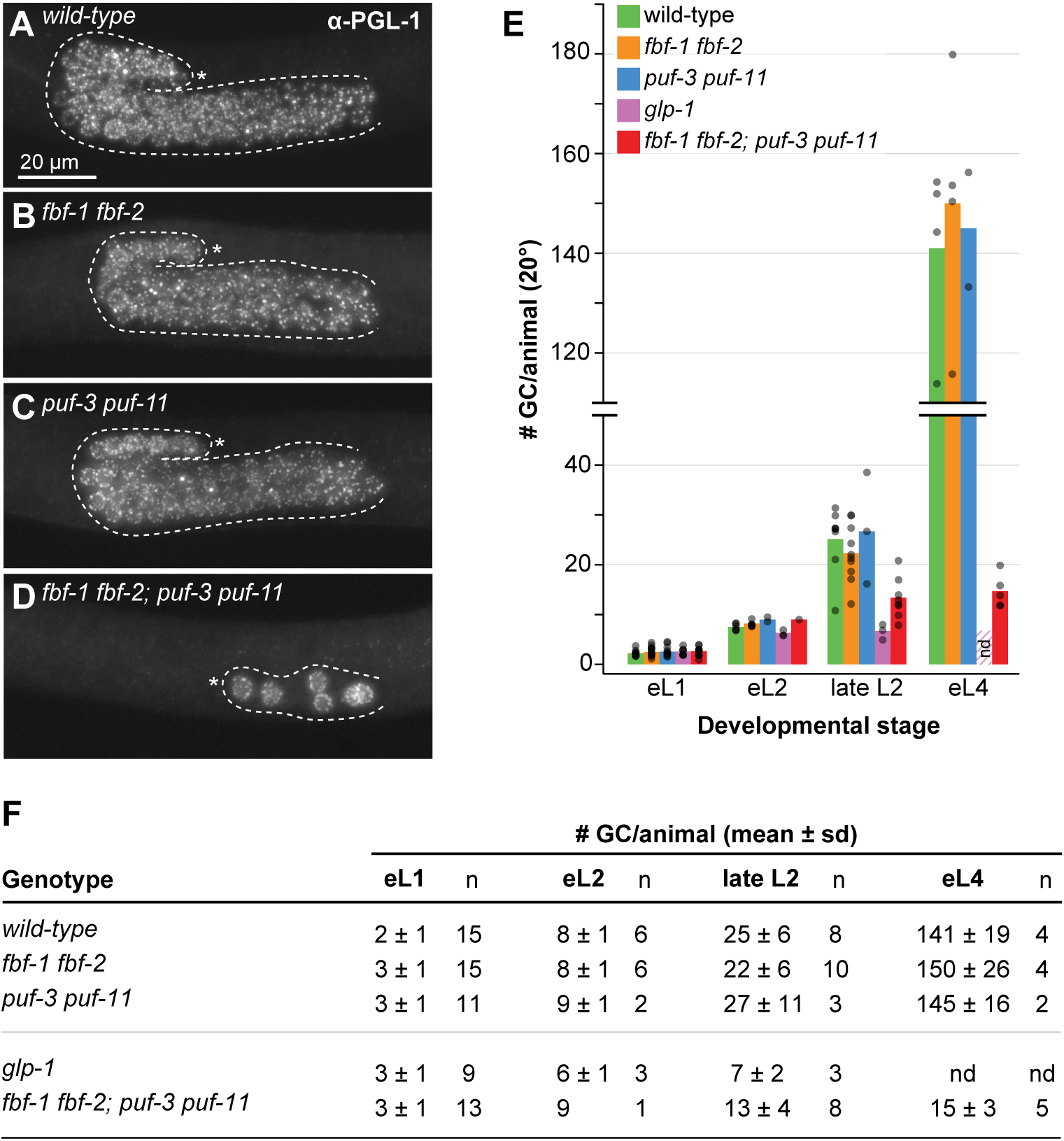
GSC self-renewal fails during larval development in *fbf-1 fbf-2; puf-3 puf-11* quadruple mutants at 20°. **A-D.** Representative images of early L4 germlines stained with α-PGL-1 (white), a germ cell marker. Images were obtained by fluorescent compound microscopy. The scale bar in A applies to all images. All other annotations by convention in Figure 1F,G. **E,F.** Quantitation of the number of germ cells (GC) per animal across larval stages. Larvae were immunostained (as in A-D) at defined developmental timepoints: early L1 (eL1), early L2 (eL2), late L2 and early L4 (eL4) stages. Germ cell number was counted as number of PGL-1-positive cells for all timepoints except early L1 was scored live by DIC microscopy. **E.** Bars show the mean number of germ cells per sample with genotypes color coded as shown. Each individual data point is plotted as a gray circle. nd, not done: germ cells in *glp-1* animals at early L4 stage had already differentiated to sperm and were thus not scored; the striped bar in early L4 is replicated from the late L2 mean value for *glp-1*. **F.** Table with mean number of germ cells (GC) per animal. **A-F.** Alleles are as follows: *fbf-1(ok91), fbf-2(q704), puf-3(q966), puf-11(q971), glp-1(q46)*.

**Figure 5:**
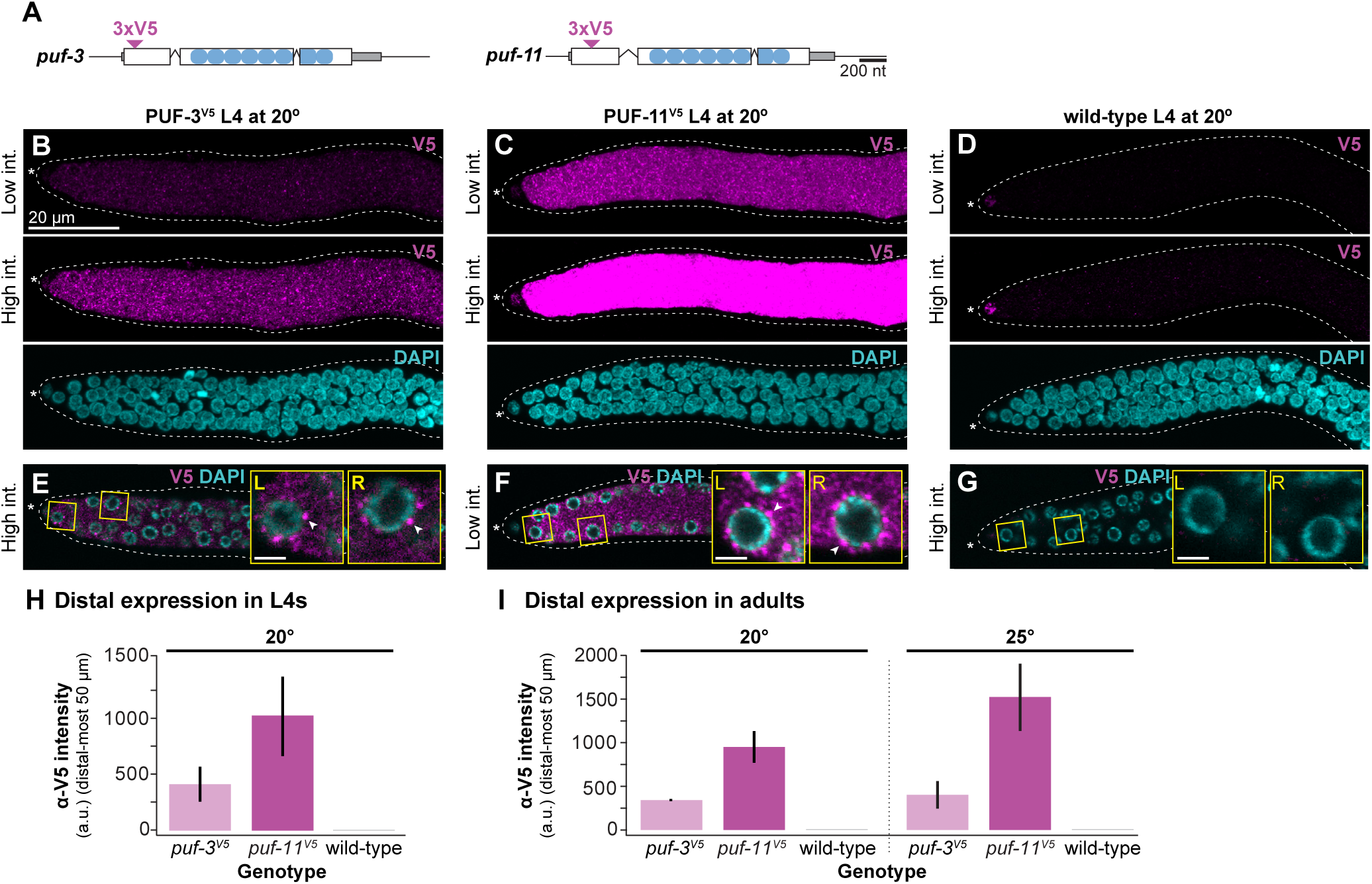
PUF-3 and PUF-11 are expressed in distal germline, including GSCs. **A.** Schematic of *puf-3* and *puf-11* loci with sites of epitope tags annotated, by convention as in Figure 1D. Inverted triangles denote insertion sites of 3xV5 epitope tags, which are flanked both up- and downstream by a GS linker. See Figure S2 for a more detailed sequence annotation. **B-G.** Representative images of PUF-3^V5^ and PUF-11^V5^ expression in gonads extruded from L4 hermaphrodites raised at 20°. Gonads from *puf-3(q1058)* (B,E), *puf-11(q1128)* (C,F) and wild-type control (D,G) were stained with α-V5 (magenta) and DAPI (cyan) and then imaged by confocal microscopy. B-D are Z-projections. V5 signal intensity (int.) was adjusted uniformly in Adobe Photoshop across images, and high or low intensities are indicated at left. E-G are single Z-slices selected from a middle plane of the same gonads imaged above; white arrowheads in insets mark representative cytoplasmic granules. Scale bar in B applies to all images except the insets, where scale bars are 2 μm. Other image annotation conventions as in Figure 1F,G. **H,I.** Quantitation of V5 intensity in the distal region of PUF-3^V5^ (light pink), PUF-11^V5^ (dark pink) and wild-type (gray) extruded gonads, determined using Fiji/ImageJ (see Methods for details). Each bar represents the mean α-V5 immunostaining intensity in arbitrary units (a.u.) in the distal-most 50 μm of the gonad (∼11 germ cell diameters (gcd) using the conversion 4.4 μm/gcd. (LEE *et al*. 2016)), with nonspecific staining background from the wild-type control subtracted. Error bars represent standard error. **H.** Analysis of mid-L4 staged extruded germlines at 20°. Four independent experiments were performed for a total of at least 27 gonads per genotype. **I.** Analysis of germlines extruded from staged adults at 20° (24 hours past mid-L4) and 25° (18 hours past mid-L4). Two independent experiments were performed for a total of 24 gonads per experimental condition. For representative images at 20°, see Figures S4E-G.

### Fluorescence quantitation

Quantitation of fluorescence in Figures 5H,I was performed with Fiji/ImageJ (SCHINDELIN *et al*. 2012). In Figure 5H, we performed four independent immunostaining experiments and quantitated a total of at least 27 gonads per genotype. In Figure 5I, we performed two independent experiments and quantitated at least 24 gonads per genotype. From confocal image stacks, we collected raw pixel intensity data from each gonad image by projecting the sum of all Z-slices onto a single plane. A freehand line, 50 pixels wide and 50 μm long that bisected the gonad, was drawn manually using the Plot Profile tool starting at the distal tip of the tissue. We found the mean intensity value of the plot profile of each gonad to get a single value reflecting the amount of protein present. Next, we subtracted any signal representing nonspecific antibody binding for each independent experiment: we calculated the mean intensity value in the respective wild-type control gonads, then subtracted it from each experimental sample. In Figure 5H,I, we report the background subtracted mean and standard error.

### Genetic epistasis experiments

To test the relationship between *lst-1 sygl-1* and *fbf-1 fbf-2; puf-3 puf-11,* we used transgenes that ubiquitously express LST-1 and SYGL-1 protein. Because these *lst-1(gf)* and *sygl-1(gf)* transgenes cause germline tumors and are sterile, they were maintained on *lst-1* or *sygl-1* RNAi, respectively, prior to the experiment. Expression of LST-1 or SYGL-1 was then induced by transferring L4s to OP50-seeded NGM plates and passaging for several generations (SHIN *et al*. 2017). The assay for Figure 6A was performed at 25°. From populations grown for 9 generations on OP50, we plated L4s onto *puf-3/11* RNAi. Next, progeny of *puf-3/11* RNAi treated animals were staged to 24 hours past L4 and stained with DAPI to assay effects on germline development. Because *lst-1* and *sygl-1* require *fbf-1 fbf-2* for function, neither *lst-1(gf)* nor *sygl-1(gf)* formed a germline tumor in these experiments.

**Figure 6:**
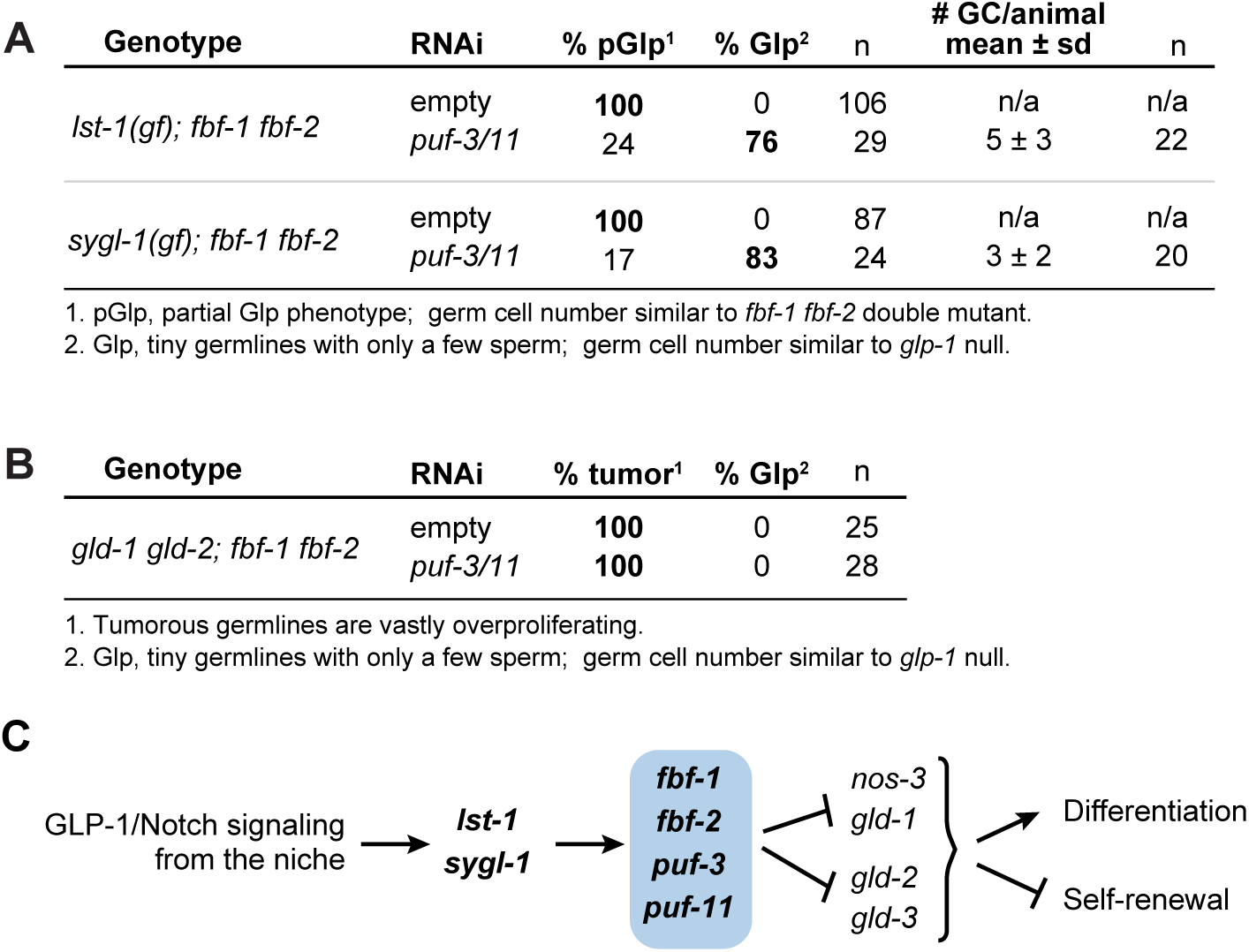
*puf-3 puf-11* lie parallel to *fbf-1 fbf-2* in the GSC regulatory pathway. **A.** Results of epistasis tests conducted with *lst-1(gf)* and *sygl-1(gf)* alleles. Number of germ cells (GC) in Glp animals was determined by counting sperm in adults and dividing by four. Genotype for *lst-1(gf)* strain is *lst-1(ok814); qSi267 [Pmex-5::LST-1::3xFLAG::tbb-2 3’ end] fbf-1(ok91) fbf-2(q704)* and *sygl-1(gf)* is *sygl-1(tm5040); qSi235 [Pmex-5::SYGL-1::3xFLAG::tbb-2 3’ end] fbf-1(ok91) fbf-2(q704*). We used the *puf-3/11* RNAi clone B in these experiments. Images of representative germlines are available in Figure S5A-D. **B.** Results of epistasis tests conducted with *gld-1 gld-2*. Genotype is *gld-2(q497) gld-1(q361); fbf-1(ok91) fbf-2(q704).* We used the *puf-3/11* RNAi clone B in these experiments. Images of representative germlines are available in Figure S5E,F. **C.** Revised pathway for GSC regulation that includes *puf-3* and *puf-11* at the same position in the pathway as *fbf-1* and *fbf-2* (blue circle).

To test the relationship between *gld-2 gld-1* and *fbf-1 fbf-2; puf-3 puf-11* for Figure 6B, we plated mid-L4 staged JK5778 to *puf-3/11* RNAi plates at 20°. F1 progeny were staged to 24 hours past L4, then DAPI stained and imaged using compound microscopy to assay germline phenotype.

### Statistical analysis

Welch’s ANOVA and Games-Howell *post hoc* tests were performed to calculate statistical significance for multiple samples. All statistical tests were performed in R and the *p*-value cut off was 0.05.

### Yeast two-hybrid

Modified yeast two-hybrid assays were performed as described (BARTEL AND FIELDS 1997). PUF proteins were fused to the LexA DNA binding domain as follows: cDNA sequences encoding PUF repeats of FBF-1 (amino acids 121-614), FBF-2 (121-632), PUF-3 (88-502), PUF-9 (162-703) and PUF-11 (91-505) were each cloned into the *Nde* I site of pBTM116 using the Gibson assembly method (GIBSON *et al*. 2009). We also used full-length LST-1 (1-328) and SYGL-1(1-206) fused to Gal4 activation domain in the pACT2 vector (SHIN *et al*. 2017). More details about plasmids are available in Table S3. To test for protein-protein interactions between PUFs and LST-1/SYGL-1, activation and binding domain pairs were co-transformed into a L40-*ura3* strain (*MATa*, *ura3-52*, *leu2-3,112*, *his3Δ200, trp1Δ1, ade2, LYS2::(LexA-op)_4_* –*HIS3*, *ura3::(LexA-op)_8_ –LacZ*) using the LiOAc method (GIETZ AND SCHIESTL 2007). *LacZ* reporter activity was measured using the Beta-Glo® Assay system (Promega #E4720), following commercial protocols and yeast-specific methods (HOOK *et al*. 2005). Luminescence was quantitated using a Biotek Synergy H4 Hybrid plate reader with Gen5 software.

### Western blot

For the western blot in Figure 7C, we grew yeast transformants in -Leu -Trp liquid media and prepared samples by boiling yeast in sample buffer (60mM Tris pH 6.8, 25% glycerol, 2% SDS, 0.1% bromophenol blue with 14 mM beta-mercaptoethanol or 100 mM DTT). Analysis was conducted on a 4-15% SDS-PAGE gradient gel (Biorad #456-1083). We probed with primary antibodies overnight at 4° as follows: 1:50,000 mouse anti-HA (HA.11, Covance #MMS-101R), 1:1000 mouse anti-V5 (1:1000, SV5-Pk1, Bio-Rad #MCA1360), or 1:10,000 mouse anti-actin (C4, Millipore #MAB1501). For secondary antibodies, blots were incubated for 1-2 hours at RT with 1:20,000 donkey anti-mouse horseradish peroxidase (Jackson ImmunoResearch #715-035-150). Immunoblots were developed using SuperSignal^TM^ West Pico/Femto Sensitivity substrate (Thermo Scientific #34080, #34095) and a Konica Minolta SRX-101A medical film processor. For final figure preparations, intensity of the blot was linearly adjusted in Adobe Photoshop.

**Figure 7:**
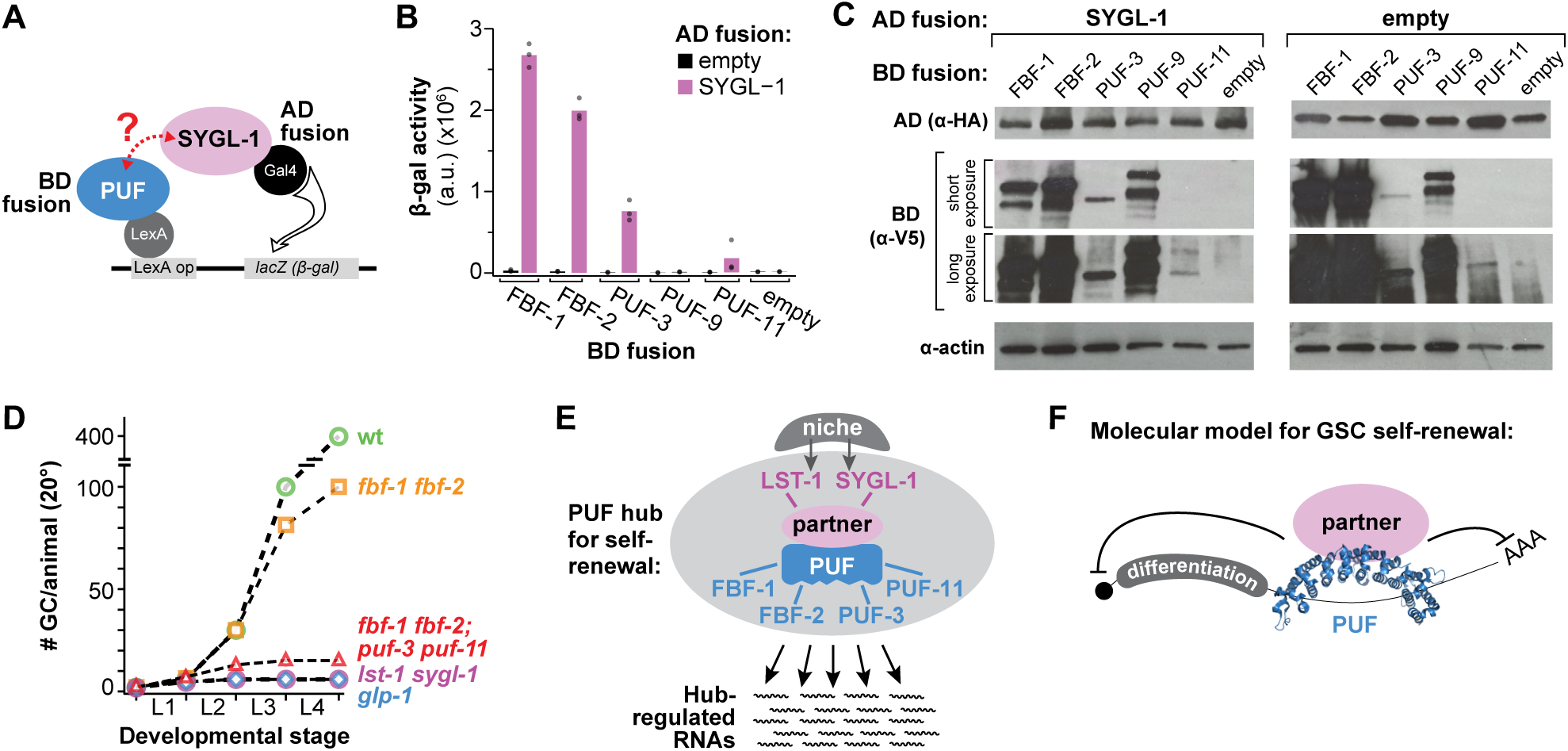
PUFs and FBFs comprise a PUF hub that accounts for GSC self-renewal. **A.** Yeast two-hybrid schematic. SYGL-1 was fused to the Gal4 activation domain (AD), which was HA tagged. PUF protein variants were fused to the LexA binding domain (BD), which was V5 tagged. PUF constructs included the PUF repeats and some flanking amino acids; for amino acid boundaries, see Methods. Interaction between SYGL-1 and PUF drives transcription of a *lacZ* (β-gal) reporter. **B.** SYGL-1/PUF interaction was measured using β-galactosidase (β-gal) activity. Each bar is the mean of at least 3 experiments. Individual data points are plotted as gray circles. **C.** Western blot from yeast lysate probed with α-V5 to detect AD fusion proteins and α- HA to detect BD fusion proteins. Actin was the loading control. **D.** Total number of germ cells (GC) per animal at 20° in mutants of key GSC regulators, revised from Figure 1B to include *fbf-1 fbf-2; puf-3 puf-11* (red triangles). **E.** The PUF hub for GSC self-renewal consists of two PUF partners that are direct targets of niche signaling and four PUF RNA-binding proteins that collectively regulate a battery of mRNAs. **F.** Molecular model for GSC self-renewal: PUF RNA-binding protein binds to the 3’ UTR of target mRNAs. A PUF partner, LST-1 or SYGL-1, ensures RNA repression by an unknown mechanism.

### Data and reagent availability

The authors affirm that all data necessary for confirming the conclusions of this article are present within the article, tables, figures, and supplemental material. All strains and plasmids are available upon request or via the Caenorhabditis Genetic Center, supported by the NIH Office of Research Infrastructure Programs (P40 OD010440). All protocols are available upon request.

## RESULTS

### Solidifying evidence for the existence of gene X

The existence of a missing GSC self-renewal regulator, gene X, was proposed because of striking differences in GSC defects upon removal of either the GLP-1/Notch receptor or its target genes (*lst-1* and *sygl-1*) on the one hand and removal of FBF-1 and FBF-2 on the other (Figure 1B) (AUSTIN AND KIMBLE 1987; CRITTENDEN *et al*. 2002; KERSHNER *et al*. 2014). To provide comprehensive data as a critical baseline for this study, we scored GSC defects in key mutants at 15°, 20° and 25° (Figure 1C). As expected, mutants lacking GLP-1/Notch or both its target genes, *lst-1* and *sygl-1*, generated only about four germ cells at each temperature, but *fbf-1 fbf-2* double mutants made many more (Figure 1C). In addition, *fbf-1 fbf-2* mutants raised at 25° possessed distal germ cells in mitotic metaphase or anaphase, consistent with active divisions (Figure S1A). A previous study showed that these distal cells express the mitotic marker, nuclear REC-8 (see SHIN *et al*. 2017, Fig S5G). Thus, loss of FBF-1 and FBF-2 has a much less severe and later effect on GSC maintenance than loss of niche signaling or loss of the niche targets, confirming the notion of some missing self-renewal regulator (gene X, Figure 1A).

### PUF-3 and PUF-11 proteins are likely the missing self-renewal regulators

To begin our search for the missing GSC regulators, we considered other PUF proteins as logical candidates. Among 10 *C. elegans* PUF-encoding genes (LIU *et al*. 2012), PUF-11 piqued our interest because of two similarities with FBF: the PUF-11 protein interacts with LST-1 in a genome-wide yeast two-hybrid screen (BOXEM *et al*. 2008), and its RNA binding specificity is similar to that of FBF (BERNSTEIN *et al*. 2005; KOH *et al*. 2009). Because the *C. elegans* genome encodes a PUF-11 paralog with nearly identical sequence, called PUF-3 (Figure 1D, Figure S2), we tested both for a role in GSC maintenance. To this end, we used two feeding RNAi clones, one targeting *puf-3* (RNAi A) and the other targeting a distinct region of *puf-11* (RNAi B) (Figure 1D). Although these RNAi clones target different gene regions, each was expected to deplete both *puf-3* and *puf-11* because of their ∼90% sequence identity (HUBSTENBERGER *et al*. 2012).

A previous *puf-3/11* RNAi study, performed in wild-type animals, identified a role for PUF-3 and PUF-11 in oogenesis, but not GSC maintenance (HUBSTENBERGER *et al*. 2012). We therefore sought a more GSC-specific assay and turned to enhancement of the *fbf-1 fbf-2* pGlp phenotype at 25°. For this analysis, we scored the presence or absence of a progenitor zone (PZ), the distal region where germ cells have not yet entered into the meiotic cell cycle and continue mitotic divisions (Figure 1E). Whereas all *fbf-1 fbf-2* adults on empty vector RNAi possessed a PZ at 25° (Figure 1F,H), most treated with *puf-3/11* RNAi lost their PZ to differentiation (Figure 1G). A comparable effect was seen in both sexes (Figure 1H). Strikingly, GSC divisions generated only ∼4 germ cells per gonad in each sex (Figure 1I), as determined by counting the number of mature sperm in adults and dividing by four. RNAi directed against other *puf* loci (e.g. *puf-8*) did not enhance the pGlp *fbf-1 fbf-2* phenotype (Figure S1B). Therefore, *puf-3/11* RNAi enhances the *fbf-1 fbf-2* germline phenotype from its partial pGlp to the full Glp typical of GLP-1/Notch mutants. This enhancement indicates that PUF-3 and PUF-11 are likely the missing self-renewal regulators.

### *puf-3* and *puf-11* mutants have no GSC proliferation defects on their own

The *puf-3/11* RNAi experiments could not distinguish between the *puf-3* and *puf-11* genes for effects on GSC maintenance. To test their individual roles, we analyzed *puf-3* and *puf-11* single mutants: three deletions, *puf-3(q801), puf-3(q966)* and *puf-11(q971),* and one nonsense allele, *puf-11(gk203683)* (Figure 2A, Figure S2). As a measure of general germline function, we scored fertility, number of embryos laid and embryo viability. The single mutants were fertile with brood sizes comparable to wild-type, and their embryos hatched into young larvae, except for *puf-11(q971)*, which had a partially penetrant embryonic lethality (Figure 2B). In addition, we scored PZ lengths as a proxy for effects on GSCs. The PZ lengths, measured with the conventional metric of germ cell diameters (gcd) from the distal end, were roughly the same as wild-type in all *puf-3* and *puf-11* single mutants (Figure 2B). Therefore, *puf-3* and *puf-11* single mutants have no major GSC proliferation defects.

We next assessed *puf-3 puf-11* double mutants (Figure 2B-F). Germlines had an organization and size comparable to wild-type (Figure 2C, D), but they made no viable embryos (Figure 2B). Male *puf-3 puf-11* crossed to feminized *fog-1* hermaphrodites yielded ample cross progeny, so *puf-3 puf-11* sperm are functional. By contrast, wild-type males crossed to *puf-3 puf-11* hermaphrodites failed to make viable progeny, suggesting that *puf-3 puf-11* oocytes are defective, consistent with prior studies (HUBSTENBERGER *et al*. 2012; HUBSTENBERGER *et al*. 2013). PZ lengths were comparable to wild-type, scored at 20° (Figure 2B), and number of germ cells therein were comparable to wild-type at both 20° and 25° (Figure 2E,F). Therefore, *puf-3 puf-11* double mutants have an oogenesis defect, as previously recognized, but no obvious GSC defects.

### *puf-3* and *puf-11* enhancement of *fbf-1 fbf-2* partial Glp phenotype

To further explore *puf-3* and *puf-11* roles in GSC maintenance, we tested for enhancement of the *fbf-1 fbf-2* pGlp phenotype using *fbf-1 fbf-2; puf* triple mutants as well as *fbf-1 fbf-2; puf-3 puf-11* quadruple mutants. To score enhancement, we determined total germ cells made in each strain by counting mature sperm number in adults and dividing by four (Figure 3A,B); at 25°, we scored for the persistence of a PZ in adults (Figure 3C,D).

We first assayed triple mutants raised at 20° (Figure 3A, left; Figure S3). The control *fbf-1 fbf-2* double mutants made roughly 100 germ cells before GSCs were lost to spermatogenesis, as previously described (CRITTENDEN *et al*. 2002; LAMONT *et al*. 2004; this work). That number decreased in both triple mutants, but the extent of pGlp enhancement differed for *puf-3* and *puf-11*. The decrease in germ cell number was small in *fbf-1 fbf-2; puf-3* triple mutants and statistically significant for only one allele; the decrease was larger in *fbf-1 fbf-2; puf-11* mutants and statistically significant for both alleles. Nonetheless, each *puf* gene contributed to larval GSC proliferation at 20°.

We also assayed triple mutants raised at 25° (Figure 3A, right; Figure 3B; Figure S3). The control *fbf-1 fbf-2* double mutants made more than 100 germ cells and maintained a PZ, as described previously (MERRITT *et al*. 2008; SHIN *et al*. 2017; this work). In *fbf-1 fbf-2; puf-3* triple mutants, germline size was comparable to *fbf-1 fbf-2* double mutants (Figure 3A, right), but many PZs were lost to meiotic entry (Figure 3B). By contrast, *fbf-1 fbf-2; puf-11* triple mutants at 25° made far fewer germ cells overall than *fbf-1 fbf-2* (Figure 3A, right), and had a fully penetrant PZ loss with all germ cells differentiating as sperm (Figure 3B). Thus, each *puf* gene enhanced the *fbf-1 fbf-2* pGlp defect at 25°, with *puf-11* again having a more severe effect than *puf-3*.

We next assayed *fbf-1 fbf-2; puf-3 puf-11* quadruple mutants, this time raised at 15°, 20° or 25°. The two distinct quadruple mutants, *fbf-1 fbf-2; puf-3(q966) puf-11(q971)* and *fbf-1 fbf-2; puf-3(q801) puf-11(gk203683)*, were remarkably similar at all three temperatures. Adults had tiny germlines composed entirely of mature sperm. Upon quantitation (Figure 3C; Figure S3), quadruples made a total of 4-9 germ cells on average at 15° and 25°, and 11-16 at 20°. This quadruple Glp phenotype is thus comparable to *glp-1* and *lst-1 sygl-1* null mutants. Thus, PUF-3 and PUF-11 function during larval development to maintain GSC divisions.

We finally asked if PUF-3 and PUF-11 maintain the PZ in *fbf-1 fbf-2* double mutant adults at 25°. To address this question, we treated mid-L4 *puf-3(q966) puf-11(q971)* double mutants with *fbf-1/2* RNAi and assayed PZ presence or absence in adults (48 hours later). Most wild-type animals treated with *fbf-1/2* RNAi retained a PZ (92%, n=12) (Figure 3D). However, few *puf-3 puf-11* double mutants treated with *fbf-1/-2* RNAi retained a PZ (7%, n=30) (Figure 3E). Instead, distal-most germ cells entered early meiotic prophase, visualized by a nuclear “crescent” morphology. Thus, PUF-3 and PUF-11 function in 25° *fbf-1 fbf-2* adults to maintain a progenitor zone.

### GSC maintenance fails during early larval development in quadruple mutants

Up to this point in this work, germ cell counts were performed in adults by counting sperm. This approach quantitates number of cells generated and differentiated, but cannot detect cells that die without differentiation into gametes. Although germ cell death was not seen in *glp-1* or *lst-1 sygl-1* mutants (AUSTIN AND KIMBLE 1987; KERSHNER *et al*. 2014), the *fbf-1 fbf-2; puf-3 puf-11* quadruple mutant may be different.

To begin, we assessed overall germline sizes at the L4 stage, which were comparable in wild-type, *fbf-1 fbf-2* and *puf-3 puf-11*, but much smaller in quadruple mutants (Figure 4A-D). Next, we counted total germ cell number at specific intervals during larval development. For this experiment, we scored *fbf-1 fbf-2; puf-3 puf-11* quadruple mutants and several controls (wild-type, *fbf-1 fbf-2* doubles*, puf-3 puf-11* doubles and *glp-1*), all maintained at 20°. PGL-1 staining was used to identify germ cells for counting (except for early L1, which was scored by DIC in live animals). We found that germ cell numbers increased similarly during larval development in wild-type, *fbf-1 fbf-2* and *puf-3 puf-11* animals, but they did not increase appreciably in *glp-1* or the *fbf-1 fbf-2; puf-3 puf-11* quadruple mutant (Figure 4E,F). By L4, the quadruple mutants had made a total of 15 germ cells on average (range=12-20, n=5) (Figure 4E,F), similar to the number of germ cells estimated from adult sperm number at the same temperature (Figure 3B,C) and consistent with cell death having little impact on germ cell number. In parallel, we visualized meiotic entry with DAPI staining. By late L2, germ cells in quadruple mutants had entered meiotic prophase, whereas wild-type, *fbf-1 fbf-2* and *puf-3 puf-11* had not. No morphological sign of germ cell death was seen over the course of these experiments. Together, these findings allay the concern that *puf-3 puf-11* mutants might reduce germ cell number by promoting cell death and thus support the conclusion that *puf-3* and *puf-11* normally promote self-renewal during larval development.

### PUF-3 and PUF-11 expression in GSCs

To test whether PUF-3 and PUF-11 proteins are expressed in GSCs, we generated V5 epitope tagged alleles (Figure 5A; Figure S2). Both PUF-3^V5^ and PUF-11^V5^ are functional, as assayed by their lack of *fbf* enhancement (Figure S4A). Upon immunostaining and imaging, PUF-3^V5^ and PUF-11^V5^ proteins were observed in the distal germlines of mid-L4 hermaphrodites raised at 20° (Figure 5B-D; Figures S4B-D for full gonad), of adult hermaphrodites raised at 20° (Figure S4E-G) and of mid-L4 males raised at 20° (Figure S4H-J). In addition, both proteins were present more proximally in developing oocytes (Figure S4E-G). As expected for PUF proteins, both PUF-3^V5^ and PUF-11^V5^ were cytoplasmic (Figure 5E-G) and localized to perinuclear granules (Figure 5E,F insets), consistent with an RNA regulatory role. No ⍺-V5 staining was seen in wild-type germlines, as expected because they lacked the epitope tag. We quantified the PUF-3^V5^ and PUF-11^V5^ signal in the distal gonads of L4s and adults at 20° (Figure 5H,I), subtracting the very low background in wild-type for each. PUF-11 was more abundant than PUF-3 at both stages. Finally, we confirmed expression of both proteins in adult distal germlines at 25° and again quantitated the signal (Figure 5I). We conclude that PUF-3 and PUF-11 are expressed in GSCs.

### *puf-3 puf-11* placement in GSC regulatory pathway

The notion that *puf-3* and *puf-11* represent the missing self-renewal regulators, dubbed gene X, predicts their placement in the GSC regulatory pathway (Figure 1A). Their *fbf* enhancement, reported above, is consistent with *puf-3 puf-11* functioning in parallel to *fbf-1 fbf-2*, but we conducted two additional epistasis experiments to solidify that pathway position. For these experiments, we used RNAi to deplete *puf-3* and *puf-11*, both for ease of genetic manipulation and because GSC defects were comparable after RNAi and in quadruple mutants.

We first investigated the relationship between *puf-3 puf-11* and *lst-1 sygl-1* (Figure 6A; Figure S6A-D). Previous studies showed that *fbf-1 fbf-2* functions either downstream or in parallel to *lst-1 sygl-1.* This pathway placement was deduced using gain-of-function (gf) mutants of *lst-1* and *sygl-1.* Both *lst-1(gf)* and *sygl-1(gf)* make massive germline tumors when *fbf-1* and *fbf-2* are wild-type, but acquire a pGlp phenotype when *fbf-1* and *fbf-*2 are removed in *lst-1(gf)*; *fbf-1 fbf-2* and *sygl-1(gf)*; *fbf-1 fbf-2* triple mutants (SHIN *et al*. 2017). To ask if *puf-3* and *puf-11* have the same genetic relationship to *lst-1* and *sygl-1*, we treated *lst-1(gf); fbf-1 fbf-2* and *sygl-1(gf); fbf-1 fbf-2* triple mutants with either empty vector RNAi as a control or *puf-3/11* RNAi. The control germlines had a pGlp phenotype (Figure 6A, Figure S6A,C), as shown previously (SHIN *et al*. 2017). However, with *puf-3/11* RNAi, most germlines had a fully Glp phenotype, typical of *fbf-1 fbf-2; puf-3 puf-11* quadruple mutants. Their germlines were tiny with only a few sperm (Figure S6B,D), and upon quantitation, only 4-8 germ cells were made on average per animal (Figure 6A). Therefore, *puf-3* and *puf-11* likely function downstream or in parallel to *lst-1* and *sygl-1* (Figure 6C).

We next investigated the relationship between *puf-3 puf-11* and *gld-1 gld-2* (Figure 6B; Figure S6E,F). The *gld-1* and *gld-2* genes promote meiotic entry and in their absence, the germline becomes tumorous (KADYK AND KIMBLE 1998). Previous studies showed that *fbf-1 fbf-2* functions upstream of *gld-1 gld-2.* Thus, *gld-1 gld-2* tumors are epistatic to *fbf-1 fbf-2* pGlp (ECKMANN *et al*. 2004). To ask if *puf-3 and puf-11* are similarly upstream of *gld-1 gld-2*, we treated *gld-1 gld-2; fbf-1 fbf-2* quadruple mutants with *puf-3/11* RNAi. Indeed, *gld-1 gld-2* tumors were still found, demonstrating *gld-1 gld-2* epistasis over *fbf-1 fbf-2; puf-3/11* RNAi (Figure 6B, Figure S6E,F). We confirmed that *puf-3,11* RNAi knockdown was successful by comparison with embryos of wild-type siblings (Figure S5G). Therefore, *puf-3* and *puf-11* function upstream of *gld-1 gld-2* (Figure 6C). Together, these experiments place *puf-3* and *puf-11* into the GSC regulatory pathway in a position consistent with their proposed identity as *gene X*.

### PUF-3 interacts with SYGL-1 in yeast

Placement of *puf-3 puf-11* alongside *fbf-1 fbf-2* in the genetic pathway (above), together with their molecular identity as PUF RNA binding proteins, suggests that PUF-3 and PUF-11 may have molecular activities in GSCs similar to FBF. Previous studies showed that FBF-1 and FBF-2 physically interact with Notch targets LST-1 and SYGL-1 (SHIN *et al*. 2017; QIU *et al*. 2019). Moreover, the LST-1 and SYGL-1 partnerships with FBF are essential *in vivo* for GSC self-renewal (HAUPT *et al*. 2019; C.R. Kanzler and H.J. Shin, unpublished). We therefore considered the possibility that PUF-3 and PUF-11 also form partnerships with LST-1 and SYGL-1. Indeed, a genome-wide interaction screen had already shown that an interaction between PUF-11 and LST-1 in yeast (BOXEM *et al*. 2008). We therefore focused our yeast two hybrid assays (Figure 7A) on interactions between PUF-3/PUF-11 and SYGL-1. FBF-1 and FBF-2 were tested as positive controls, and PUF-9 as a likely negative control. PUF-3 interacted robustly with SYGL-1, but PUF-9 and PUF-11 did not (Figure 7B). The discrepancy between PUF-3 and PUF-11 was initially confounding given the similarity of the two proteins. However, a Western blot revealed that PUF-11 was poorly expressed in yeast (Figure 7C). We conclude that PUF-3 interacts with SYGL-1 and suggest that PUF-11 likely does as well, due to the near identity between PUF-3 and PUF-11. These data plus those of the genome-wide screen (BOXEM *et al*. 2008) indicate that PUF-3 and PUF-11 likely interact physically with LST-1 and SYGL-1.

## DISCUSSION

### Missing self-renewal regulators are PUF-3 and PUF-11

Despite over three decades of research defining the regulatory network that maintains *C. elegans* germline stem cells, key self-renewal regulators were clearly missing (see Introduction, Figure 1A,B). The argument was simple. Animals lacking GLP-1/Notch signaling from the niche or lacking the two key GLP-1/Notch targets, *lst-1* and *sygl-1,* have a much earlier GSC defect than animals lacking the only other known self-renewal regulators, FBF-1 and FBF-2. Because LST-1 and SYGL-1 proteins interact physically with FBF-1 and FBF-2 (SHIN *et al*. 2017; HAUPT *et al*. 2019; QIU *et al*. 2019) and that interaction is essential for GSC self-renewal (HAUPT *et al*. 2019), we predicted that the missing regulators might be additional PUF RNA-binding proteins. Indeed, we report here that animals lacking four PUF proteins, PUF-3 and PUF-11 in addition to FBF-1 and FBF-2, exhibit the same early GSC defect as *glp-1* null mutants (Figure 7D). The *fbf-1 fbf-2; puf-3 puf-11* germ cell number is equivalent to that of *glp-1* null at 15° and 25°. At 20°, the quadruple mutant undergoes one additional GSC division, which is a minor difference (one temperature, one cell division). Thus, PUF-3 and PUF-11 are the major missing self-renewal regulators.

In the process of characterizing *puf-3* and *puf-11* as self-renewal regulators, we confirmed their primary role in oogenesis. Previous studies using RNAi directed against *puf-3/11* had identified their function in oogenesis – namely, to produce viable embryos (HUBSTENBERGER *et al*. 2012). Our work extends that previous study in three ways. Using deletion mutants of each gene, we find that PUF-3 and PUF-11 act redundantly during oogenesis; using tagged versions of each protein, we show that both are expressed in oocytes; and using genetics, we find that PUF-3 and PUF-11 are not required for spermatogenesis. Importantly, while their oogenesis function is exclusive to hermaphrodites, their role in GSC self-renewal is critical in both hermaphrodites and males and thus is gender-independent. Their molecular mechanism of action in both GSCs and oocytes likely revolves around RNA regulation, a common theme among PUF proteins. Regardless, we emphasize that PUF-3 and PUF-11 are the long-sought missing self-renewal regulators.

### A “PUF hub” is responsible for GSC self-renewal

Regulatory networks are central to cell fates and tissue patterning across animal phylogeny. For germline fates and patterning, regulatory networks that rely on post-transcriptional regulation have emerged as particularly prominent (e.g. KIMBLE AND CRITTENDEN 2007; SLAIDINA AND LEHMANN 2014; YAMAJI *et al*. 2017), with key RNA-binding proteins regulating hundreds of RNAs and directing cell fate programs (KERSHNER AND KIMBLE 2010; AOKI *et al*. 2018; PORTER *et al*. 2019; this work). This work reveals that PUF RNA-binding proteins are principle intrinsic regulators in the stem cell network. Four PUFs collectively drive GSC self-renewal in hermaphrodites and males, in larvae and adults and at all laboratory growth temperatures (15°, 20° and 25°). The centrality of PUF proteins to the GSC regulatory network is underscored by the fact that GSC defects of *fbf-1 fbf-2; puf-3 puf-11* quadruple null mutants are virtually identical to those of *glp-1*/Notch null and *lst-1 sygl-1* null mutants (Figure 7D).

The remarkable phenotypic congruence of mutants in niche signaling, its targets LST-1 and SYGL-1, and four PUF proteins leads us to propose the concept of a “PUF self-renewal hub” in the stem cell regulatory network (Figure 7E). This hub consists of four PUF RNA-binding proteins and two PUF partner proteins, LST-1 and SYGL-1. LST-1 and SYGL-1 were first identified as partners of FBF (SHIN *et al*. 2017; HAUPT *et al*. 2019; QIU *et al*. 2019), but PUF-3 and PUF-11 also likely partner with LST-1 and SYGL-1 (BOXEM *et al*. 2008; this work). Moreover, these PUF partnerships are essential for GSC self-renewal (HAUPT *et al*. 2019; C.R. Kanzler and H.J. Shin, unpublished). The PUF hub therefore serves as the principal node for GSC self-renewal in the stem cell regulatory network

The PUF hub seems remarkably simple and is strongly supported by genetic and molecular analyses. However, puzzles remain. For example, mutants in the *fog-1* gene, which encodes a CPEB-related RNA-binding protein, also enhance the GSC defect of *fbf-1 fbf-2* double mutants, but that enhancement is coupled to a reversal in germline sex and the mechanism remains a mystery (THOMPSON *et al*. 2005). In contrast, as emphasized here, the “PUF hub” GSC phenotype is not coupled to any effect on germline sex determination, but instead is equivalent to removal of niche signaling (KERSHNER *et al*. 2014; this work). Most other intrinsic stem cell regulators do not meet this high bar of equivalence to the niche-defective phenotype. Thus, the PUF hub promises to provide a paradigm for understanding self-renewal hubs more broadly.

### Redundancy and buffering within the PUF hub

The PUF hub relies on a striking nexus of functional redundancies. PUF-3 and PUF-11 are redundant with FBF during larval development and in adults at 25° (this work); and FBF-1 and FBF-2 are redundant with each other in late larval development at 15° and 20° (CRITTENDEN *et al*. 2002). Moreover, the two PUF partners, LST-1 and SYGL-1, are functionally redundant (KERSHNER *et al*. 2014; SHIN *et al*. 2017). These layers of redundancy, together with our molecular understanding of individual hub proteins, suggests a simple molecular model (Figure 7F). In this model, each PUF protein binds to target RNAs via 3’UTR regulatory elements and also binds to either LST-1 or SYGL-1 to elicit RNA repression. Evidence for this model is particularly strong for the FBFs, whose mode of action has been analyzed most intensively (BERNSTEIN *et al*. 2005; WANG *et al*. 2009; SHIN *et al*. 2017; HAUPT *et al*. 2019). Data are also strongly suggestive for the nearly identical PUF-3 and PUF-11 proteins: PUF-11 binds to RNA with a sequence specificity similar to that of FBF (KOH *et al*. 2009); PUF-3/-11 repress expression of reporter RNAs in oocytes (HUBSTENBERGER *et al*. 2012); PUF-11 interacts with LST-1 in yeast (BOXEM *et al*. 2008); and PUF-3 interacts with SYGL-1 in yeast (this work). While further analyses are needed, outlines of the hub architecture and definition of its key molecular features are clear.

The extensive redundancy among hub components suggests that components are, to a first approximation, molecularly interchangeable. That interchangeability likely renders the hub robust, namely capable of maintaining stem cells under many conditions (e.g. developmental stage, growth temperature, sex). Although not yet analyzed, the interchangeability may also help stem cells withstand the barrage of environmental inputs and stresses experienced outside the laboratory. A similar phenomenon of functional redundancy of key regulators has been found in other developmental regulatory networks (e.g. Hox genes in animal development, MAD box genes in plant development) and likely lies at the heart of network evolution more broadly (WAGNER 2008).

In addition to functional redundancy, single hub components likely have specialized individual roles. Intensive studies of FBF-1 and FBF-2 reveal numerous individual features (LAMONT *et al*. 2004; VORONINA *et al*. 2012; VORONINA 2013; BRENNER AND SCHEDL 2016; PRASAD *et al*. 2016; WANG *et al*. 2016; PORTER *et al*. 2019). FBF-1 and FBF-2 have distinct low penetrance sex determination defects, genetic interactions, expression, subcellular localization, target RNAs and FBF-specific molecular effects on targets. PUF-3 and PUF-11 will also likely possess differences, between each other and also with the FBFs. Understanding the common and unique roles among the members of the hub will be crucial to understanding how the hub is buffered to maintain stem cells under a variety of physiological and environmental conditions.

## ACKNOWLEDGEMENTS

We thank Peggy Kroll-Conner for help generating strains central to this work as well as isolation of the *puf-3(q801)* deletion, Maureen Barr for sharing her mutagenesis library, and the Million Mutations Project for generating *puf-11(gk203683).* We thank Susan Strome (University of California, Santa Cruz) for α-PGL-1 and α-SP56 antibodies. We are also grateful to Sarah Crittenden and Brian Carrick for comments on the manuscript, Laura Vanderploeg for help with figures and Carol Pfeffer for help with manuscript preparation. Some strains used in the study were provided by the *Caenorhabditis* Genetics Center, supported by the NIH Office of Research Infrastructure Programs (P40 OD010440).

