## Supplementary Figures and Tables for "A PUF hub drives self-renewal in *C. elegans* germline stem cells"

**Supplementary Figure 1**
**Haupt et al**

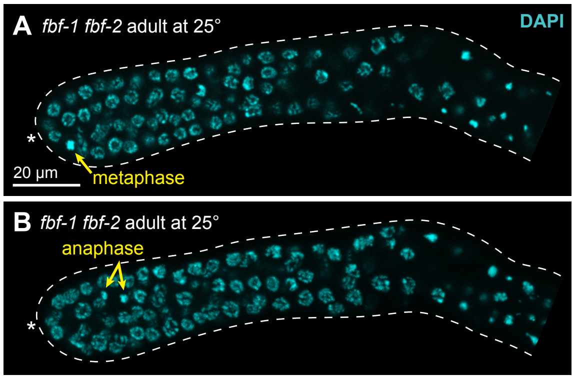


**C RNAi against other *pufs***

| Genotype | RNAi | Distal germ cells, adults at 25° % PZ | % sperm | n |
|---|---|---|---|---|
| *fbf-1 fbf-2* | empty | **87** | 0 | 12 |
| | *puf-5* | **93** | 0 | 14 |
| | *puf-6* | **66** | 0 | 22 |
| | *puf-7* | **72** | 0 | 16 |
| | *puf-8* | **86** | 0 | 18 |
| | *puf-9* | **100** | 0 | 17 |

### Figure S1: *fbf-1 fbf-2* mutant characterization

**A, B**. Representative images of gonads extruded from *fbf-1 fbf-2* adult hermaphrodites that had been staged to 18 hours after L4 at 25° and stained with DAPI (cyan). Morphologies consistent with metaphase (A) and anaphase (B) are annotated (yellow arrows). Other conventions as in Figure 1F,G. The images in A and B show distinct confocal Z-sections from the same gonad. The scale bar in A applies to both images. Alleles used here and throughout are *fbf-1(ok91)* and *fbf-2(q704)*.

**C**. State of distal germ cells in *fbf-1 fbf-2* adult hermaphrodites raised at 25° on either empty RNAi or RNAi against various *puf* genes. All were staged to 18 hours after L4. Germ cell states were scored as described in legend to Figure 1H.

```
q966 begins               q1058 (GS linker::3xV5::GS linker insertion)      q801 begins
PUF-3  MSQNTGSSNLGRYYESPPTATEAR-----GTFGGCFNANSSTNIWTPNRKVDSSMGFRSSSPTPQSAWAPNRQGFGGQFSKWRTSTPMTTPARYPQQAVRLID   98
PUF-11 MSQSTGSSNLGRYHESPPT-TEARNVSGKNTFGGCFNSNSS-NIWTPNRNVDSSMGFQRVSSTPKNASTPYRQGFGGQFSKWRTSTPMTTPARHPQQALRLID  101
q971 begins                  q1128 (GS linker::3xV5::GS linker insertion)

PUF-3  LENNASFSKSLNSTTRSHKCTLPIWAGDGEGNVSDSVTLQDVLANDALVEFATDKNGCRFLQEHYPTENDNDVHQKLFRKLVEDRAIFLSLCSNMFGNFFVQR  201
PUF-11 LENNASSPNTLNSSTRSYKCTLPIWAGDGEGNVSDSVTLPDVLANDALVEFATDKNGCRFLQEHYPTESDNDIHQKLFRKLVEDRAIFLSLCCNMFGNFFVQR  204
                                                                   gk203683 (R160Opal)
                                                                                                   q801 ends
PUF-3  VLECSNTEEQEILTEHLATDLYNLCLDKSACRVIQLAIQKLDVHLATRLSLELRDTHLVRLSIDQNGNHVIQKIVKTLPVSSWTFLVDFFADDDNLIHVCQDK  304
PUF-11 VLECSNTEEQEILTEHLASDLYNLCLDKSACRVIQLAIQKLDVHLATRLSLELRDTYLVRLSIDQNGNHVIQKIVKTLPVSAWSFVVEFFADDDNLIHVCQDK  307

PUF-3  YGCRVIQSTVETLSTDQYAQCYQHRVILLRSLMAGVTRNCTQLASNEFANYVVQHVIKCGDALAVYRDIIIEQCLLQNLLSMSQEKYASHVVEVAFECAPYRL  407
PUF-11 YGCRVIQSTVETLSSDTYAECYQQRVVLLRSLMSGVTRNCSQLASNEFANYVVQHVIKCGDAMAVYRDVIIEQCLLQNLLSMSQEKYASHVVEVAFGCAPCRL  410
                                                                                               q966 ends
PUF-3  VAEMMNEIFEGYIPHPDTNRDALDILLFHQYGNYVVQQMIQTCVLGQNARDQKQSEMYGMWLEKIHGRVMRNAHRLERFSSGKKIIEALQSMSLY          502
PUF-11 AAEMMNEIFEGYIPHPDTNRDALDILLFHQYGNYVVQKMIQICVLGQNARDQKQAEMYGMWLEKIRERVMRNANRLERFSSGKKIIEALQSMSFY          505
                                                                                               q971 ends
```

**Figure S2: Annotated PUF-3 and PUF-11 protein sequences**

Alignment of PUF-3 and PUF-11 amino acid sequences to show: molecular endpoints of key deletion mutants *q966, q801* and *q971* (black brackets); nonsense mutant *gk203683* (red); insertion sites for GS linker::3xV5::GS linker in *puf-3*$^{V5}$*(q1058)* and *puf-11*$^{V5}$*(q1128)* (magenta triangles); residues in PUF repeats (blue) (Zhang *et al.* 1997), with amino acids critical for RNA interaction (bold) (Wang *et al.* 2002).

| | # GC/animal (mean ± sd) | | | | | | | | |
|---|---|---|---|---|---|---|---|---|---|
| **Genotype** | **15°** | n | *p*-value | **20°** | n | *p*-value | **25°** | n | *p*-value |
| *fbf-1 fbf-2* | 91 ± 20 | 16 | n/a | 128 ± 30 | 20 | n/a | many & still dividing | 50 | n/a |
| *fbf-1 fbf-2; puf-3(q966)* | nd | nd | nd | 84 ± 19 | 10 | 0.002 | many & meitoic entry | 43 | n/a |
| *fbf-1 fbf-2; puf-3(q801)* | nd | nd | nd | 102 ± 13 | 10 | 0.061 | many & meitoic entry | 50 | n/a |
| *fbf-1 fbf-2; puf-11(q971)* | nd | nd | nd | 25 ± 9 | 10 | <0.001 | 11 ± 7 | 20 | n/a |
| *fbf-1 fbf-2; puf-11(gk203683)* | nd | nd | nd | 42 ± 11 | 20 | <0.001 | 14 ± 8 | 20 | n/a |
| *fbf-1 fbf-2; puf-3(q966) puf-11(q971)* | 7 ± 3 | 20 | <0.001 | 16 ± 7 | 20 | <0.001 | 4 ± 2 | 20 | n/a |
| *fbf-1 fbf-2; puf-3(q801) puf-11(gk203683)* | 5 ± 4 | 23 | <0.001 | 11 ± 8 | 11 | <0.001 | 9 ± 5 | 8 | n/a |

**Figure S3: Triple and quadruple mutant germ cell counts**

Number of germ cells (GC) made in *fbf-1 fbf-2; puf* triple mutants and *fbf-1 fbf-2; puf-3 puf-11* quadruple mutants at 15°, 20° and 25°. Germlines were scored as described in Figure 1C, with the following modification: GC number were not counted in larger germlines with distal germ cells in meiotic prophase but not yet sperm; these were scored as “many & meiotic entry”. nd, not done. *p*-value compared to respective *fbf-1 fbf-2* control was determined using Welch’s ANOVA and Games-Howell *post-hoc* test; n/a, not applicable.

**A**

| Genotype | Distal germ cell state, adults at 25° | | | |
|---|---|---|---|---|
| | % PZ | % meiotic | % sperm | n (arms) |
| *fbf-1 fbf-2* | **97** | 3 | 0 | 100 |
| *fbf-1 fbf-2; puf-3$^{V5}$(q1058)* | **96** | 4 | 0 | 100 |
| *fbf-1 fbf-2; puf-11$^{V5}$(q1128)* | **99** | 1 | 0 | 100 |

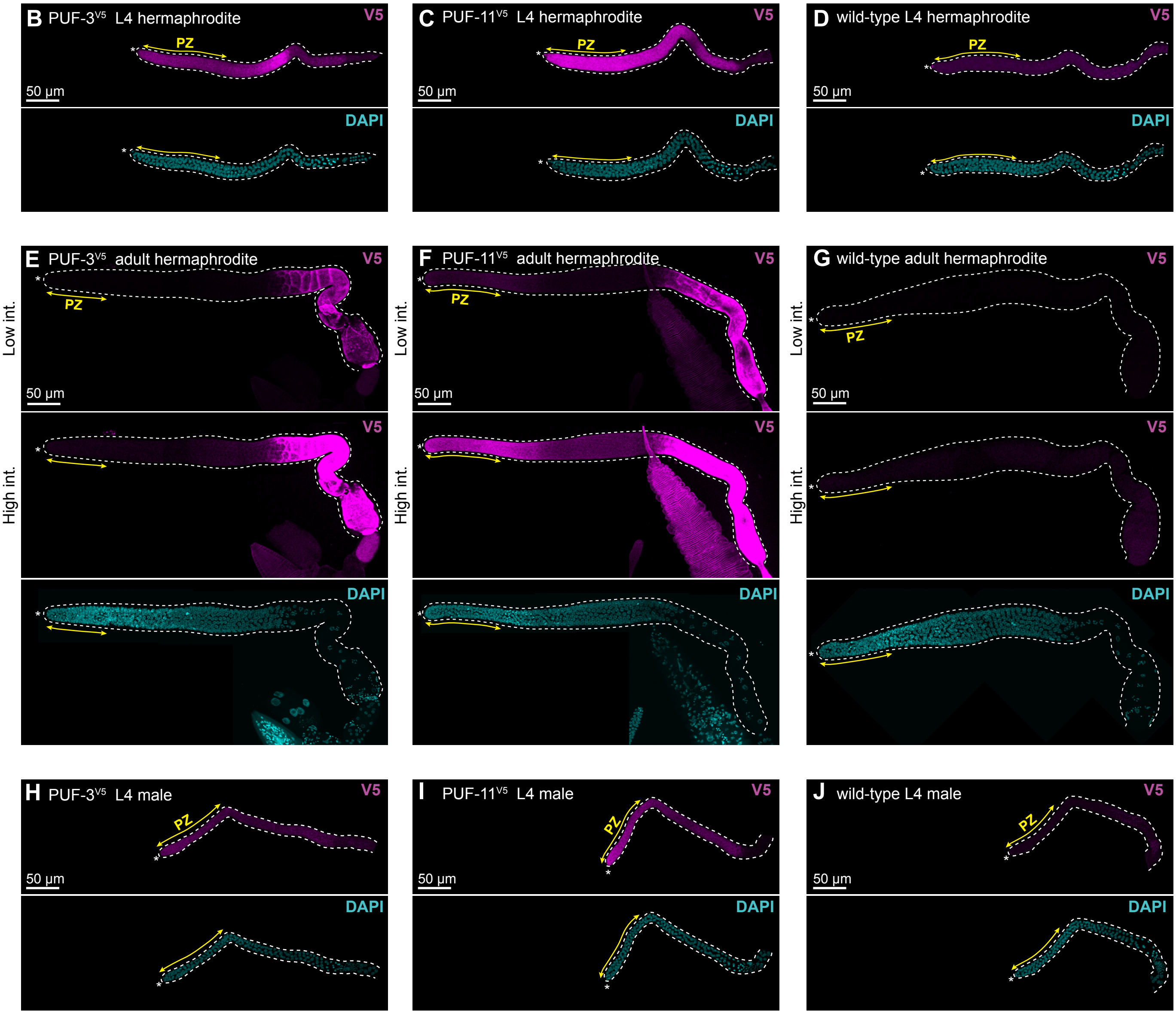


**Figure S4: PUF-3$^{V5}$ and PUF-11$^{V5}$ function and expression in whole gonads**

**A.** Epitope-tagged alleles of *puf-3* and *puf-11* did not enhance pGlp phenotype of *fbf-1 fbf-2* mutants and hence made functional protein. Strains were raised at 25° and assayed 18 hours past L4. Germ cell state was scored as described in Figure 1H and 3B.

**B-J.** Representative images of PUF-3$^{V5}$ and PUF-11$^{V5}$ expression in whole gonads at 20°. Gonads were extruded from *puf-3(q1058)* (B,E,H), *puf-11(q1128)* (C,F,I) and wild-type control (D,G,J) animals and then stained with α-V5 (magenta) and DAPI (cyan). Images are maximum intensity Z projections taken by confocal microscopy. PZ is indicated by a double headed arrow (yellow), other annotation convention as in Figure 1F,G.

**B-D.** Gonads from mid-L4 staged hermaphrodites.

**E-G.** Gonads from adult hermaphrodites staged to 24 hours past mid-L4. Intensity (int.) of the V5 signal was adjusted uniformly across images in Adobe Photoshop, with high or low intensity indicated at left.

**H-J.** Germlines from mid-L4 staged males.

**Supplementary Figure 5**
**Haupt et al**

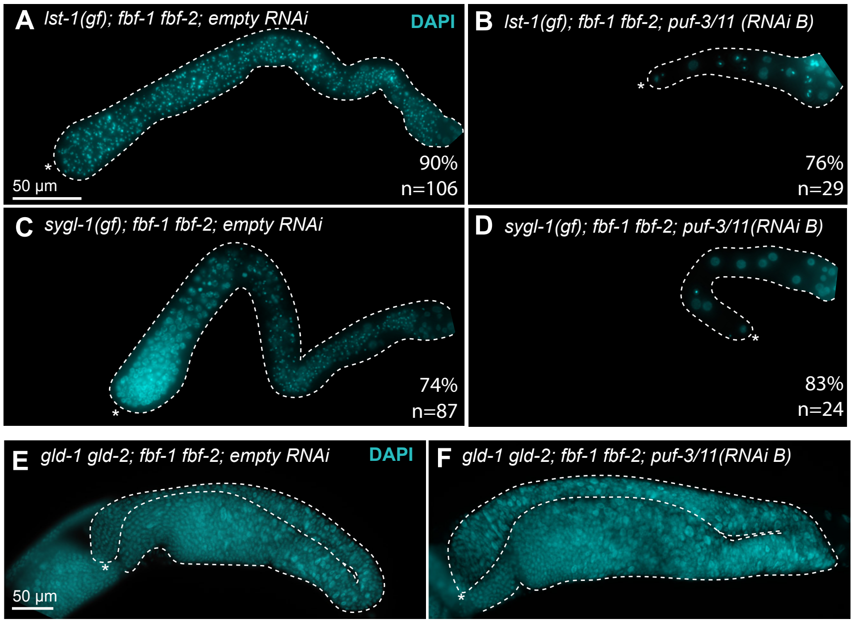


**G**

| **Genotype** | **RNAi** | **% animals with abnormal embryos *in utero*** | n |
|---|---|---|---|
| *gld-1 gld-2/+; fbf-1 fbf-2/+* | empty | 1 | 79 |
| | *puf-3/11* | 100 | 74 |

### Figure S5: Epistasis results

**A-D.** Epistasis tests using *lst-1(gf)* and *sygl-1(gf)*. Representative compound microscopy images of gonads extruded from adults staged to 18 hours past L4 at 25°, stained with DAPI (cyan) and then scored for the Glp phenotype (4-8 germ cells differentiated as sperm, as defined in Austin and Kimble (1987)). We expected undifferentiated cells in the distal gonad on the empty RNAi, typical of *fbf-1 fbf-2* mutants at this temperature (Figure 1F). However, the distal state of differentiation was variable (also noted in Shin *et al.* 2017), so penetrance of the depicted phenotype is given at bottom right. Most *sygl-1(gf); fbf-1 fbf-2* germlines on empty RNAi had undifferentiated cells at the distal end (74%, n=87) (C), but some had differentiated. For *lst-1(gf); fbf-1 fbf-2,* only 10% remained undifferentiated while 90% had differentiated (n=106) (A). The scale bar in A applies to all images, and all conventions are as in Figure 1F,G. Genotype for *lst-1(gf)* strain is *lst-1(ok814); qSi267 [Pmex-5::LST-1::3xFLAG::tbb-2 3' end] fbf-1(ok91) fbf-2(q704)* and *sygl-1(gf)* is *sygl-1(tm5040); qSi235 [Pmex-5::SYGL-1::3xFLAG::tbb-2 3' end] fbf-1(ok91) fbf-2(q704)*. We utilized *puf-3/11* RNAi clone B in these experiments.
**E,F.** Epistasis tests using *gld-1 gld-2*. Representative compound microscopy images of adults staged to 24 hours past L4 at 20°, stained with DAPI (cyan), and scored for either a tumorous germline typical of *gld-1 gld-2* or a Glp germline (containing 4-8 germ cells differentiated as sperm, as defined in Austin and Kimble (1987). The scale bar in E applies to both images, with other conventions as in Figure 1F,G. Genotype is *gld-2(q497) gld-1(q361); fbf-1(ok91) fbf-2(q704).* We used *puf-3/11* RNAi clone B in these experiments.
**G.** Confirmation that *puf-3/11* RNAi worked in *gld-1 gld-2* epistasis test. Effective *puf-3/11* RNAi causes embryo lethality (Hubstenberger *et al.* 2012; this work). We therefore DAPI stained and scored for abnormal morphology of *in utero* embryos in *gld-1 gld-2/+* siblings to show that *puf-3/11* had been successfully knocked down.

**S1 Table. Nematode strains used in this study**

| Name | Genotype | Reference |
|---|---|---|
| N2 | wild-type | Brenner, 1974 |
| EG7866 | *oxTi564 II; unc-119(ed3) III* | Frøkjær-Jensen, 2014 |
| JK3107 | *fbf-1(ok91) fbf-2(q704)/ mIn1[mIs14 dpy-10(e128)] II* | Crittenden, 2002 |
| JK3108 | *fbf-1(ok91) fbf-2(q704)/ mIn1[mIs14 dpy-10(e128)] II; him-5(e1490) V* | this work |
| JK3743 | *fog-1(q785) I/ hT2[qIs48](I;III)* | Thompson, 2005 |
| JK4256 | *puf-3(q801) IV/ nT1[qIs51(IV;V)* | this work |
| JK4862 | *glp-1(q46) III/ hT2[qIs48](I;III)* | Kershner, 2014 |
| JK5411 | *sygl-1(tm5040) I; fbf-1(ok91) fbf-2(q704) qSi235/ mIn1[mIs14 dpy-10(e128)] II* | Shin, 2017 |
| JK5537 | *lst-1(ok814) I; fbf-1(ok91) fbf-2(q704) qSi267/ mIn1[mIs14 dpy-10(e128)] II* | Shin, 2017 |
| JK5778 | *gld-2(q497) gld-1(q361)/ ccIs4251 unc-15(e73) I; fbf-1(ok91) fbf-2(q704)/ mIn1[mIs14 dpy-10(e128)] II* | this work |
| JK5908 | *puf-11(gk203683) IV* | this work |
| JK5915 | *puf-3(q966) IV* | this work |
| JK5925 | *fbf-1(ok91) fbf-2(q704)/ mIn1[mIs14 dpy-10(e128)] II; puf-11(gk203683) IV* | this work |
| JK5926 | *fbf-1(ok91) fbf-2(q704)/ mIn1[mIs14 dpy-10(e128)] II; puf-3(q966) IV* | this work |
| JK5991 | *puf-3(q801) puf-11(gk203683) IV/ nT1[qIs51] IV;V* | this work |

| | | |
|---|---|---|
| JK5993 | *fbf-1(ok91) fbf-2(q704)/ mIn1[mIs14 dpy-10(e128)] II; puf-3(q801) IV* | this work |
| JK5996 | *puf-11(q971) IV* | this work |
| JK6051 | *fbf-1(ok91) fbf-2(q704)/ mIn1[mIs14 dpy-10(e128)] II; puf-11(q971) IV* | this work |
| JK6080 | *puf-3(q1058) IV* | this work |
| JK6143 | *fbf-1(ok91) fbf-2(q704)/ dpy-10(q1074) oxTi564 II; puf-3(q801) puf-11(gk203683) IV/ nT1[qIs51](IV;V)* | this work |
| JK6272 | *fbf-1(ok91) fbf-2(q704)/ mIn1[mIs14 dpy-10(e128)] II; puf-3(q1058) IV* | this work |
| JK6280 | *puf-11(q1128) IV* | this work |
| JK6282 | *fbf-1(ok91) fbf-2(q704)/ mIn1[mIs14 dpy-10(e128)] II; puf-11(q1128) IV* | this work |
| JK6321 | *puf-3(q966) puf-11(q971) IV/ nT1[qIs51](IV;V)* | this work |
| JK6333 | *fbf-1(ok91) fbf-2(q704)/ dpy-10(q1074) oxTi564 II; puf-3(q966) puf-11(q971) IV/ nT1[qIs51](IV;V)* | this work |
| JK6401 | *lst-1(q869) sygl-1(q828) I/ hT2[qIs48](I;III)* | Haupt, 2019 |

**S2 Table. CRISPR alleles generated in this study**

| Allele | Description | Guide(s) (5'–3') | Repair template (5'–3')[1] | Parent strain |
|---|---|---|---|---|
| *q966* | *puf-3(null)* | UGCUUUGACUCAUAUUGGUA and GCCUAUGUGUAUCUAGUACA | gatgtcttccaaaaaaataaattttcaggtagtctTcgtaccaatatacacataggcaatttttattcatttccatttgaatcctaaccc | wild-type |
| *q971* | *puf-11(null)* | UGCUUUGACUCAUAUUGGUA and AAUGUCCUUUUACUAGAUAU | accatttttcccaaaaaaatattttcagttagtctAcgtaccaatatatagAcaattcatttgcattattatcctaacccccactcacgt | wild-type |
| *q1058* | *puf-3* exon 1 insertion of GS linker:: 3xV5:: GS linker | ACGGAGGCUCGUGGCACAUU | cgagagcccgccaacggcgacggaggctcgtGGATCTGGTAAGCCTATCCCTAACCCTCTCCTCGGTCTAGATAGTACTGGAAAGCCAATCCCAAACCCACTCCTCGGACTTGATAGCACCGGTAAGCCTATCCCTAACCCACTCCTCGGACTTGATAGCACCGGATCTggAacCttcggtggttgcttcaacgccaaca | wild-type |
| *q1074* | *dpy-10* frameshift[2] | GCUACCAUAGGCACCACGAG[3] | n/a | EG7866 |
| *q1128* | *puf-11* exon 1 insertion of GS linker:: 3xV5:: GS linker | UCCGGAAAAAACACAUUCGG | tccgccaacgacggaggctcgcaatgtgtccGGATCTGGTAAGCCTATCCCTAACCCTCTCCTCGGTCTAGATAGTACTGGAAAGCCAATCCCAAACCCACTCCTCGGACTTGATAGCACCGGTAAGCCTATCCCTAACCCACTCCTCGGACTTGATAGCACCGGATCTggaaaGaaTacCttTggcggttgcttcaact | wild-type |

[1] Uppercase letters denote mutations (including insertions, PAM mutations and/or seed sequence mutations)

[2] *dpy-10(wild-type):* tggaaaccgtaccgct**CG**tggtgcctatggtag
*dpy-10(q1074)*: tggaaaccgtaccgct**GCC**tggtgcctatggtag

[3] Arribere, et al. (2014)

**S3 Table. Yeast-two hybrid plasmids used in this study**

| Plasmid | Insert description | Cloning site | Vector backbone | Reference |
|---|---|---|---|---|
| pJK1580 | HA::SYGL-1(aa 1-206) | *Nco* I | pACT2 | Shin, 2017 |
| pJK2015 | HA::LST-1(aa 1-328) | *Xho* I | pACT2 | Shin, 2017 |
| pJK2033 | V5::FBF-1(aa 121-614) | *Nde* I | pBTM116 | this work |
| pJK2034 | V5::PUF-3(aa 88-502) | *Nde* I | pBTM116 | this work |
| pJK2037 | V5::PUF-11(aa 91-505) | *Nde* I | pBTM116 | this work |
| pJK2046 | V5::FBF-2(aa 121-632) | *Nde* I | pBTM116 | this work |
| pJK2056 | V5::PUF-9(aa 162-704) | *Nde* I | pBTM116 | this work |
